# Stroke-related gene expression profiles across species: A meta-analysis

**DOI:** 10.1101/2023.03.13.532480

**Authors:** Ruslan Rust

## Abstract

Stroke patients are often left with permanent disabilities with no regenerative treatment options. Unbiased RNA sequencing studies decoding the transcriptional signature of stroked tissue hold promise to identify new potential targets and pathways directed to improve treatment for stroke patients. Here, gene expression profiles of stroked tissue across different time points, species, and stroke models were compared using NCBI GEO database. In total, 32 datasets from mice, rats, humans, and primates were included, exploring gene expression differences in healthy and stroked brain tissue. Distinct changes in gene expression and pathway enrichment revealed the heterogenicity of the stroke pathology in stroke-related pathways e.g., inflammatory responses, vascular repair, remodelling and cell proliferation and adhesion but also in diverse general, stroke-unrelated pathways that have to be carefully considered when evaluating new promising therapeutic targets.

## Background

Stroke is a major cause for disability and death affecting one in four people during their lifetime [1]. Stroke survivors have only limited therapeutic options and are often left with considerable disabilities [2,3]. The development of new therapeutics for stroke is not trivial as the ischemic cascade involves many pathological processes such as cellular excitotoxicity, oxidative stress, inflammation, blood-brain barrier disruption and scarring [4]. Therefore, unbiased screening studies decoding the transcriptional response of stroked tissue are important to identify new potential targets for therapy [5,6]. Since human stroke brain tissue is only of limited suitability for the analysis (i.e., it can only be accessed in lethal strokes and the processing time may lead to poor RNA quality), the majority of transcriptional profiling of stroked tissue was performed in rodents. Although animal models are designed to generate reproducible infarcts in highly controlled conditions, there is recognized heterogeneity among stroke models and species [7]. The most commonly used stroke model is the middle cerebral artery occlusion (MCAo) model; in this model the most commonly affected artery in human stroke is surgically obscured. The occlusion can either be transient to produce reperfusion after 30-120 min (tMCAo) or permanent (pMCAo) [8,9]. Alternatively, the photothrombotic stroke model is popular for permanent ischemia in defined brain regions that lead to long-term functional deficits [9–11].

In this study, the gene expression omnibus (GEO) RNA sequencing data were obtained from stroke tissue across different stroke models, sex, time points, and species. The gene expression profiles were integrated to identify and compare common and differentially expressed genes (DEG) and enriched pathways in the individual groups.

## Methods

### Meta analysis procedure

data search from GEO microarray data repositories were searched in November 2022. The search terms were “stroke”, “ischemia”, “tMCAo”, “pMCAo”, “MCAo”, “photothrombotic stroke”. Selected organisms were “homo sapiens”, “macaca”, “mus musculus” and “rattus norvegicus”. Datasets were excluded that did not include brain tissue samples (**Suppl. Fig. 3**). Further studies were excluded with missing information, duplicates, pooled samples, poor quality controls, and no clear separation between stroked and non-stroked groups. An overview of the used datasets can be found in **Suppl. Table 1.** Significant genes for each group were identified using R Studio RankProd [12]. All datasets were annotated and converted uniformly using genome wide annotation resources.

### Study selection

The search initially retrieved 338 articles, of which 32 met the inclusion criteria [list studies].

### Functional enrichment analysis of DEGs

Analysis of RNA sequencing data and functional enrichment analysis for DEGs was performed using EdgeR [13] and clusterProfiler 4.0 [14] in RStudio.

## Results

### Gene expression profiles of stroked brain tissue across sex, species, stroke model, and time

In total, 338 studies were screened that examined gene-expression differences in stroke. Of these, 213 studies analyzed brain tissue and used non-stroked brain tissue controls. Datasets with missing information, duplicates, pooled samples, poor quality controls, and no clear separation between stroked and non-stroked groups were further excluded. In total, we included 32 datasets from mice (12), rats (15), humans (2) and primates (3) (**Fig. 1A**). The datasets included in this meta-analysis compared gene expression in stroked brain tissue to control brain tissue (either from intact, sham operated or contralesional brain regions). Detailed information of each dataset is provided in **Suppl. Table 1** describing the GEO ID, sample type, control type, stroke model, time point, sex, and further information.

**Fig. 1:**
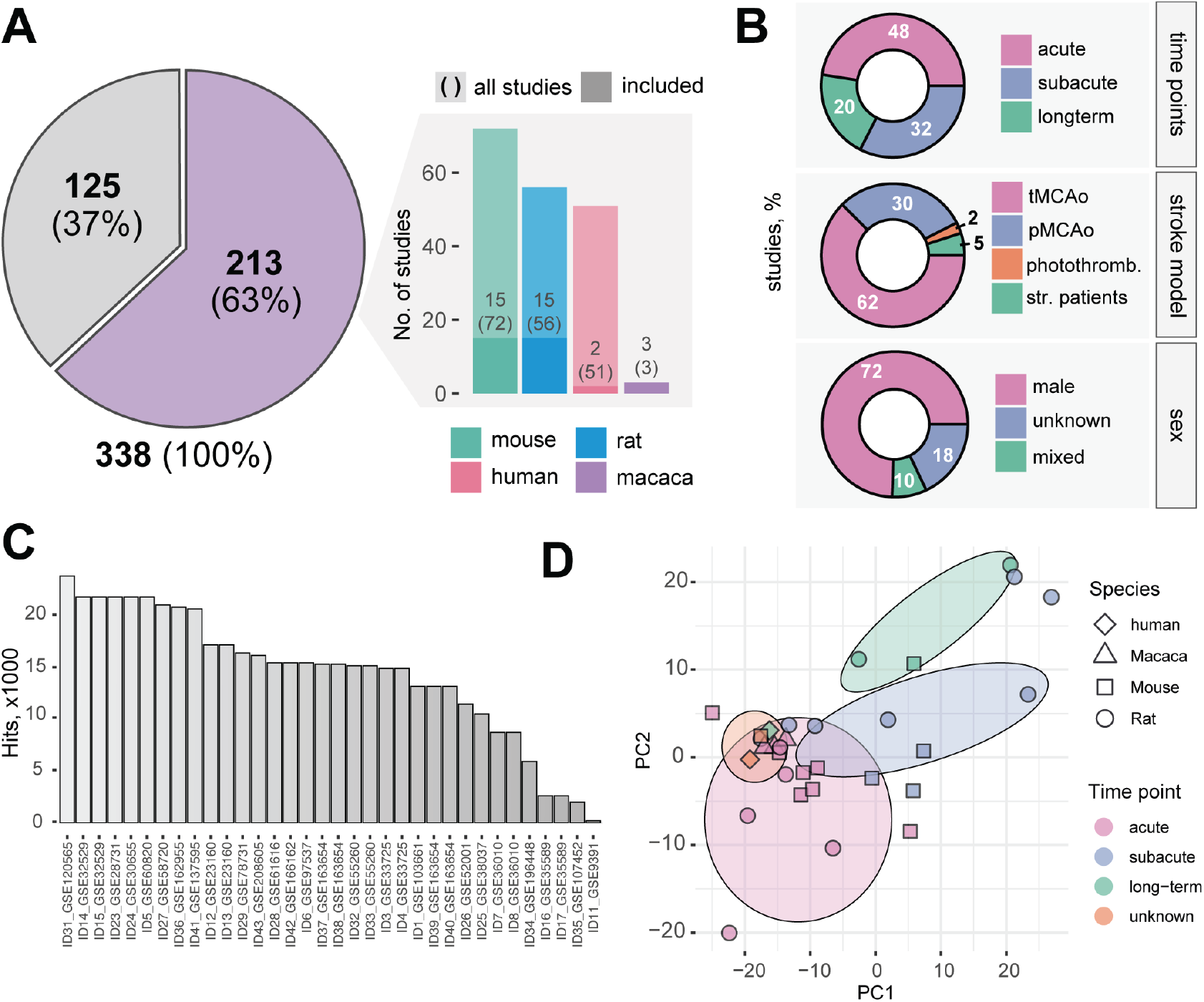
Overview of studies dissecting stroke-related gene expression differences in the brain. **(A)** (left) Excluded and included stroke studies from NCBI GEO. (Right) Exclusion of datasets subdivided by species (mouse, rat, human and primates) with missing information, duplicates, pooled samples, poor quality controls, and no clear separation between stroked and non-stroked groups were further excluded. **(B)** Distribution of individual studies by time point, stroke model and sex. **(C)** Number of gene hits in individual studies. More information about the IDs can be found in **Suppl Table 1**. **(D)** Principal component analysis (PCA) of all stroke studies across different species and time points.

The majority of datasets were acquired either in the acute, (<24h: 49%) or subacute (>24h and <6d: 36%) phase after stroke, only few datasets investigated long-term gene expression changes after stroke (>6d: 10%). The most frequently applied model of stroke was the tMCAo (67%), followed by the pMCAo model (31%) only 2% of the studies used the photothrombotic stroke model. Surprisingly, the vast majority of datasets only used the male sex (74%) (**Fig. 1B**). The majority of datasets had data from 10-25k genes, only four micro-array studies showed data for less than 10k hits (all of them were derived from human and primate samples) (**Fig. 1C**). Principal component analysis (PCA) revealed the widest separation across the datasets for the acute, subacute, and long-term periods after stroke indicating the importance of time in the transcriptomic signatures (**Fig. 1D**).

### Gene expression differences in stroked brain tissue between mice and rats are pronounced in acute phase after injury

The vast majority of stroke studies are performed in either rats or mice, hence the focus of the subsequent analysis was on the more detailed rodent datasets.

To analyze the effect of the species, the transcriptomes were compared at the acute, subacute, and long-term timepoints after MCAo stroke induction. Analysis of differentially expressed genes (DEGs) revealed that there was a greater overlap between downregulated genes in the mouse and rat transcriptome to all time points (**Fig 2 A, Suppl. Fig. 1**). The overlap was also higher at the subacute and long-term time point (shared downregulated genes: acute: 26%, subacute: 24%, long-term: 25%; shared upregulated genes: acute: 1%, subacute: 12%, long-term: 5%, **Fig. 2A**). A list of the 60 most DEG can be found in **Suppl. Tables 2-4**. Gene ontology (GO) analysis was carried out to identify the biological function of the DEGs in the mouse and rat stroke transcriptome. Interestingly, most significantly enriched GO terms were distinct at the different time points. While only 10% of the top30 GO terms in the acute phase were stroke-related, the majority of enriched GO terms in the subacute and long-term time point were directly related to the stroke pathology. For instance, inflammation related GO terms (e.g., immune system response, regulation of leukocyte activation) were differentially enriched in the subacute and long-term phase between mouse and rat. In the long-term, differences in the synaptic signaling and synaptic organization were among the top enriched GO terms (**Fig. 2B**, **Suppl. Table 8-10**).

**Fig. 2:**
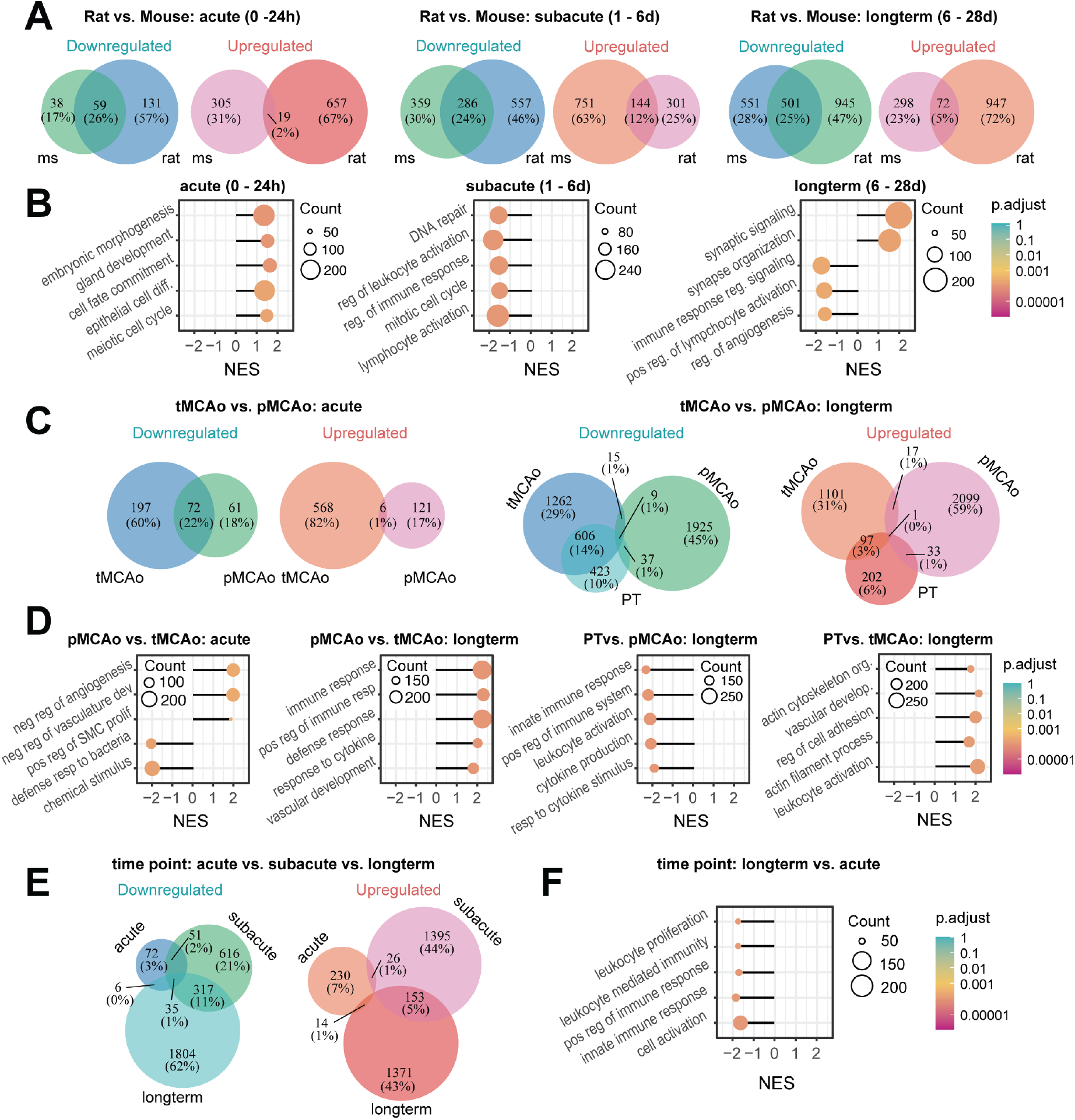
Gene expression profiles after stroke differ across species. **(A)** Venn Diagram representing number of shared and differentially expressed genes between mice and rats to different time points. **(B)** Gene ontology analysis of a subset of most significantly enriched biological processes in rats and mice. **(C)** Venn Diagram representing number of shared and differentially expressed genes between permanent MCAo and transient MCAo and the photothrombotic stroke model at different time points. **(D)** Gene ontology analysis of a subset of most significantly enriched biological processes in different stroke models. **(E)** Venn Diagram representing number of shared and differentially expressed genes between different time periods after stroke. **(F)** Gene ontology analysis of a subset of most significantly enriched biological processes at different time points.

### Different stroke-models have distinct gene expression profiles in stroked brain tissuex

Acutely after stroke transient MCAo (tMCAo) and permanent MCAo (pMCAo) are the most suitable models to mimic acute human stroke cascade [7]. These methods enable either permanent occlusion of the blood vessels or transient ischemia with reperfusion.

First gene expression datasets from stroked tissue were compared in rodents using tMCAo to pMCAo procedure at the acute (<24h) time period. The total number of up- and downregulated DEG was higher following tMCAo acutely after stroke. While 22% of common genes were downregulated in both models only 1% of common genes were upregulated acutely after stroke indicating a unique stroke signature between tMCAo and pMCAo models (**Fig. 2C, Suppl. Table 5**). Most significantly enriched pathways in tMCAo included detection of chemical stimulus and innate defense responses whereas most enriched pathway in pMCAo included e.g. regulators of vascular and smooth muscle cell responses (**Fig. 2D, Suppl. Table 11**).

Apart from MCAo models, long-term recovery after stroke can additionally be evaluated using a photothrombotic stroke model (PT). Although the PT model does cause a vasogenic edema acutely after stroke (that is uncharacteristic for human stroke), the method is minimally invasive and results in well-characterized long-term sensory-motor deficits and gradual incomplete recovery [10,15]. DEG were compared in stroked tissue of pMCAo, tMCAo and PT models at long-term (6-28 d) period. More genes were differentially expressed in pMCAo and tMCAo at a long-term period compared to the acute phase after stroke. A higher overlap could be observed for downregulated and upregulated genes (18%) in tMCAo and PT model, whereas long-term pMCAo gene expression had a highly unique molecular signature compared to the other stroke models (tMCAo: 3%; PT: 3%) (**Fig 2C, Suppl. Table 6**). Most significantly enriched pathways in the pMCAo model compared to tMCAo and PT were related to regulators of immune responses e.g., positive regulation of immune system responses, regulation of cytokine production, leukocyte activation indicating a considerably stronger immune activity in pMCAo model. Apart from immune system related pathways, the PT stroke model showed an enrichment in remodeling related pathways including, regulation of cell adhesion, actin cytoskeleton organization and tissue and blood vessel morphogenesis (**Fig 2D Suppl. Table 12-14**).

### Stroke gene expression signature is unique in the acute, subacute, and long-term post-injury time periods

As the highest deviation in the transcriptome signature after stroke appears to be dependent on the time point after stroke (**Fig. 2E, Fig. 1D, Suppl. Table 7**), I investigated which pathways were enriched to the different time points.

As expected, most immune-related pathways were strongly downregulated at the long-term time point compared to the acute phase after stroke such as leukocyte proliferation and positive regulation to immune response (**Fig. 2F, Suppl. Fig. 2).** However, in the subacute phase many processes involving synapse activation and signaling as well as neurotransmitter secretion were upregulated compared to the acute phase. Interestingly, this activation seem appears to be transient as those pathways e.g., synaptic signaling and synaptic plasticity were downregulated again at the long-term phase after stroke (**Fig 2F, Suppl. Fig 2, Suppl Table 15-17**).

## Discussion

Dissecting the molecular profile after stroke promises to identify new targets for potential therapeutic compounds. However, the multifaceted ischemic cascade and the variety of used animal models complicates the search of promising pathways related to stroke. Here, major differences were identified in publicly available gene expression profiles after stroke that varied depending on the time period, the animal/rodent species, and the stroke model used. These alterations affected primarily upregulated genes and affected both general and stroke-related pathways. The biggest alteration in the gene expression was identified for the different time periods after stroke, supporting the hypothesis that timing of therapy in stroke is of primary importance [16].

Furthermore, RNA datasets from primates and humans were only comparable to a limited extent with the rodent datasets. The human and primate datasets had considerably less analysed genes as they were derived from micro-array studies. Additionally, obtaining human stroked brain tissue is challenging as it can only be collected post-mortem.

The heterogenicity of the stroke pathophysiology is a further limitation in identifying novel targets for therapy. For instance, the molecular signature of the stroke core may considerably vary from the penumbra and different cell types may contribute to gene expression changes. However, there are only a limited number of studies that have thoroughly investigated these differences [17]. Novel advancements with single cell/nucleus and spatially resolved transcriptomics may provide further insights in near future [18,19].

This meta-analysis reveals that considerable gene expression differences exist between mice and rats as well as the used stroke models. This data supports the recommendations to confirm the effect of experimental therapeutic compounds in at least two animal species or different stroke models [20]. Additionally, the study was designed to examine the effect of sex on gene expression after stroke. However, there is still a lack of datasets for the female sex after stroke as all studies used male rodents or mixed sex. The sex plays an important role in human stroke pathology [21] and should be further investigated in preclinical gene expression studies in near future.

## Conclusion

In sum, this meta-analysis identifies distinct gene expression changes in stroked brain tissue across species, time points and stroke models. These differences affected general and stroke-related pathways and need to be considered when evaluating new potential therapeutic compounds.

## Supporting information

All Supplementary Data in xlsx format

All Supplementary Data in docx format

## Declarations

## Acknowledgement

Not applicable

## Consent for publication

The author consents to the publication of the manuscript.

## Availability of Data and Materials

All raw data are available in the supplementary files of the manuscript.

## Authors contribution

R.R. designed the study, carried out the systematic review, generated, and analyzed data, wrote and revised the manuscript.

## Ethics approval and consent to participate

Not applicable

