## Supplementary Figures for "Stroke-related gene expression profiles across species: A meta-analysis"

### Title

Ruslan Rust^1,2^

### Affiliation

^1^ Institute for Regenerative Medicine, University of Zurich, 8952 Schlieren, Switzerland,

^2^ Neuroscience Center Zurich, University of Zurich and ETH Zurich, Zurich, Switzerland

### Correspondence

Ruslan Rust
Institute for Regenerative Medicine (IREM)
University of Zurich, Campus Schlieren
Wagistrasse 12
8952 Schlieren / Zurich, Switzerland
, +41 44 63 7688

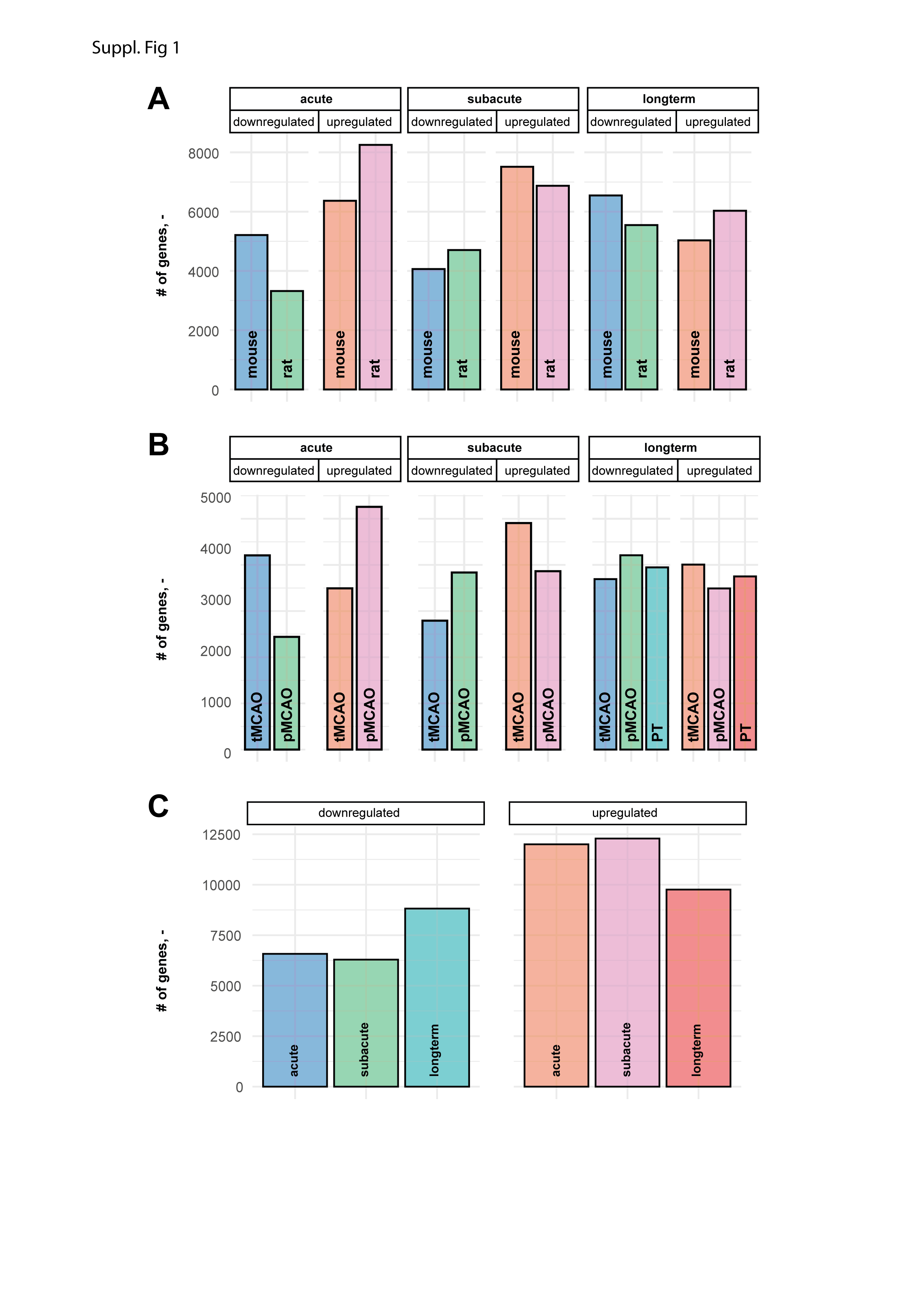

**Suppl. Fig 1: Gene expression profile after stroke.** Ratio of up- and downregulated genes in stroked brain tissue across (A) species (B) stroke model and (C) time period.

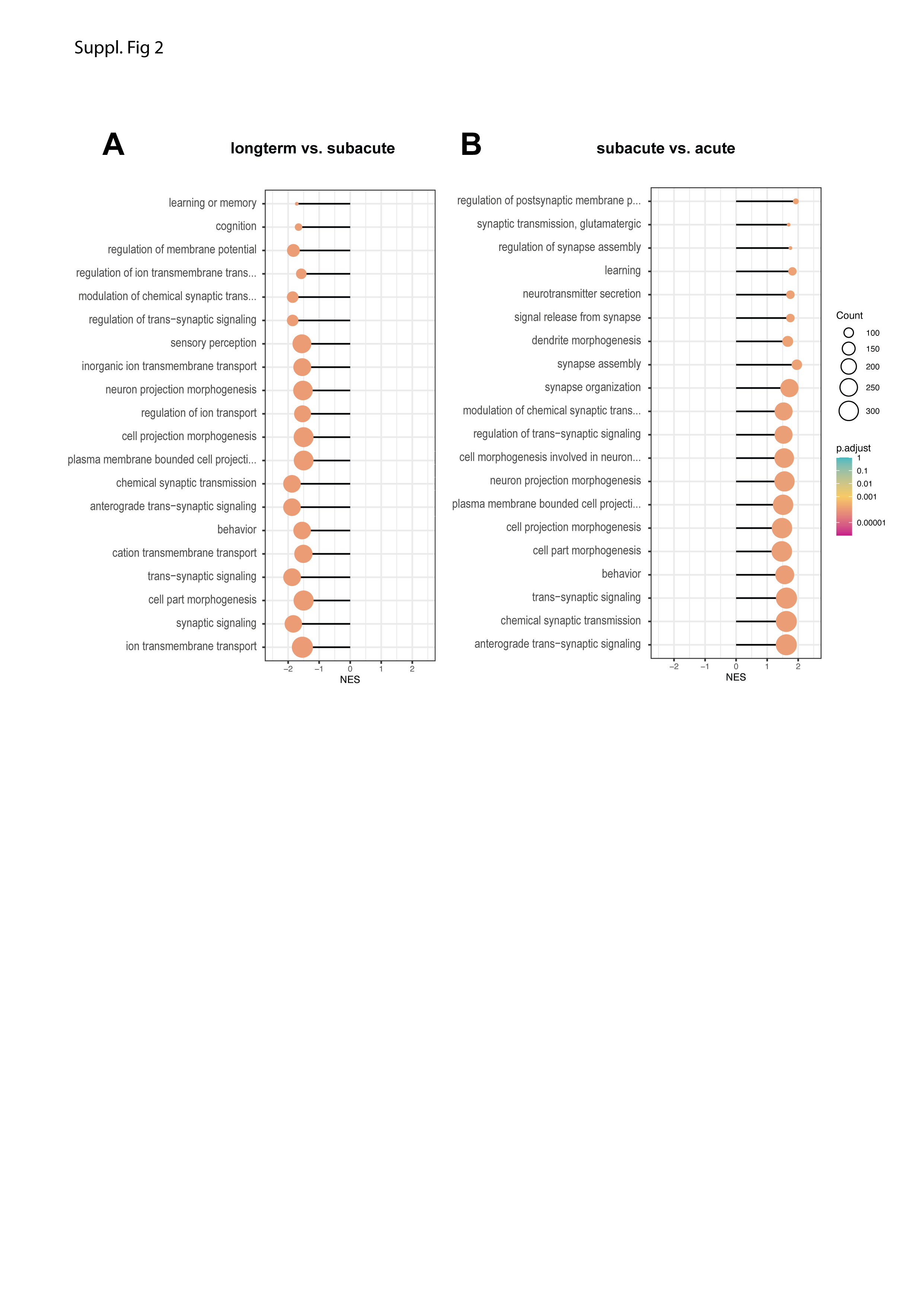

**Suppl. Fig. 2: Gene set enrichment of most enriched biological processes at different time points.** Gene ontology analysis of top20 enriched pathways between (A) long-term and subacute time period and (B) between subacute and acute time period.

**Identification of studies via databases and registers**

Records removed *before screening*:

Duplicate records removed (n = 20)

Records removed for other reasons (n = 105)

Records identified from*:

NCBI GEO (n = 338)

**Identification**

Reports excluded:

No brain tissue (n = 120)

No control group (n = 16)

Missing data (n = 42)

Reports assessed for eligibility

(n = 213)

**Screening**

Studies included

(n = 35)

**Included**

**Suppl. Fig. 3: PRISMA flow diagram for search criteria**
