## Supplementary material for "Stroke-related gene expression profiles across species: A meta-analysis": All Supplementary Data in docx format

**Suppl. Table 1: Overview of all analyzed studies from GEO database**

| **ID** | **Species** | **Sex** | **age** | **ExactTime** | **Time points** | **Stroke model** | **Reperfusion, min** | **Control** |
| --- | --- | --- | --- | --- | --- | --- | --- | --- |
| GSE103661 | Rattus norvegicus | male | adult | 3h | acute | pMCAO | permanent | sham operated |
| GSE33725 | Rattus norvegicus | male | adult | 6h | acute | pMCAO | permanent | sham operated |
| GSE33725 | Rattus norvegicus | male | adult | 2h | acute | pMCAO | permanent | sham operated |
| GSE60820 | mus musculus | male | adult | 6h | acute | pMCAO | permanent | contralesional site |
| GSE97537 | Rattus norvegicus | male | adult | 24h | subacute | pMCAO | permanent | sham operated |
| GSE36010 | Rattus norvegicus | unknown | adult | 24h | subacute | pMCAO | permanent | sham operated |
| GSE36010 | Rattus norvegicus | unknown | adult | 3d | long-term | pMCAO | permanent | sham operated |
| GSE9391 | homo sapiens | mixed | adult | different | long-term | natural | NA | intact |
| GSE23160 | mus musculus | male | adult | 8h | acute | tMCAO | 120 | sham operated |
| GSE23160 | mus musculus | male | adult | 24h | subacute | tMCAO | 120 | sham operated |
| GSE32529 | mus musculus | male | adult | 24h | subacute | tMCAO | 45 | sham operated |
| GSE32529 | mus musculus | male | adult | 3h | acute | tMCAO | 45 | sham operated |
| GSE35589 | macaca mulatta | male | adult | 1h | acute | tMCAO | 90 | contralesional site |
| GSE35589 | macaca mulatta | male | adult | 6h | acute | tMCAO | 90 | contralesional site |
| GSE28731 | mus musculus | male | adult | 24h | acute | tMCAO | 60 | sham operated |
| GSE30655 | mus musculus | male | adult | 24h | subacute | tMCAO | 60 | intact |
| GSE38037 | Rattus Norvegicus | male | adult | 24h | subacute | tMCAO | unknown | sham operated |
| GSE52001 | Rattus Norvegicus | male | adult | 6d | long-term | tMCAO | 60 | unknown |
| GSE58720 | mus musculus | male | adult | 24h | subacute | tMCAO | 90 | unknown |
| GSE61616 | Rattus Norvegicus | male | adult | 7d | long-term | tMCAO | unknown | unknown |
| GSE78731 | Rattus Norvegicus | male | adult | 72h | subacute | tMCAO | unknown | intact |
| GSE120565 | mus musculus | mixed | adult | 48h | subacute | tMCAO | 60 | contralesional site |
| GSE55260 | Rattus Norvegicus | male | adult | 14d | long-term | tMCAO | 90 | intact |
| GSE55260 | Rattus Norvegicus | male | adult | 3d | subacute | tMCAO | 90 | intact |
| GSE196448 | mus musculus | unknown | adult | 7d | long-term | tMCAO | 7 | sham operated |
| GSE107452 | macaca mulatta | male | adult | 6h | acute | tMCAO | unknown | contralesional site |
| GSE162955 | homo sapiens | mixed | adult | different | long-term | natural | NA | contralesional site |
| GSE163654 | mus musculus | unknown | adult | 2h | acute | pMCAO | permanent | sham operated |
| GSE163654 | mus musculus | unknown | adult | 6h | acute | pMCAO | permanent | sham operated |
| GSE163654 | Rattus Norvegicus | unknown | adult | 2h | acute | pMCAO | permanent | sham operated |
| GSE163654 | Rattus Norvegicus | unknown | adult | 6h | acute | pMCAO | permanent | sham operated |
| GSE137595 | mus musculus | male | adult | 2h | acute | tMCAO | 90 | sham operated |
| GSE166162 | Rattus Norvegicus | male | adult | 3d | subacute | tMCAO | 60 | sham operated |
| GSE208605 | mus musculus | mixed | adult | 21d | long-term | photothrombotic stroke | permanent | sham operated |

**Suppl. Table 2:** Median fold changes of most differentially expressed genes acutely after stroke in mice and rats

| **Species** | **Gene.symbol** | **logFC** | **Regulation** |
| --- | --- | --- | --- |
| Mouse | or5m9 | 0.819437 | Upregulated |
| Mouse | or4e5 | 0.803961 | Upregulated |
| Mouse | or1r1 | 0.639548 | Upregulated |
| Mouse | or10j27 | 0.634558 | Upregulated |
| Mouse | or5w19 | 0.633792 | Upregulated |
| Mouse | sostdc1 | 0.633737 | Upregulated |
| Mouse | or8k22 | 0.598463 | Upregulated |
| Mouse | or9k2 | 0.596986 | Upregulated |
| Mouse | or6aa1 | 0.595239 | Upregulated |
| Mouse | or52n3 | 0.589649 | Upregulated |
| Mouse | or1j13 | 0.586391 | Upregulated |
| Mouse | or52s1 | 0.584994 | Upregulated |
| Mouse | or2b2 | 0.574185 | Upregulated |
| Mouse | amy2a3 | 0.572381 | Upregulated |
| Mouse | or51q1 | 0.547148 | Upregulated |
| Mouse | or4b1 | 0.538302 | Upregulated |
| Mouse | or6d15 | 0.534399 | Upregulated |
| Mouse | or52ab2 | 0.524342 | Upregulated |
| Mouse | or10d3 | 0.512509 | Upregulated |
| Mouse | or52h1 | 0.505126 | Upregulated |
| Mouse | or2n1e | 0.504454 | Upregulated |
| Mouse | drd2 | 0.496306 | Upregulated |
| Mouse | or10d4b | 0.495224 | Upregulated |
| Mouse | or5p4 | 0.49445 | Upregulated |
| Mouse | or5w18 | 0.490622 | Upregulated |
| Mouse | or7g21 | 0.489712 | Upregulated |
| Mouse | or2y15 | 0.4883 | Upregulated |
| Mouse | kcnj13 | 0.487332 | Upregulated |
| Mouse | or14a257 | 0.48316 | Upregulated |
| Mouse | or4p22 | 0.482062 | Upregulated |
| Rat | lbp | 2.003615 | Upregulated |
| Rat | fgf10 | 1.661478 | Upregulated |
| Rat | mpl | 1.661034 | Upregulated |
| Rat | svs5 | 1.63815 | Upregulated |
| Rat | rhox12 | 1.593303 | Upregulated |
| Rat | glra3 | 1.584856 | Upregulated |
| Rat | lhx1 | 1.579322 | Upregulated |
| Rat | slc26a2 | 1.56773 | Upregulated |
| Rat | irs1 | 1.566479 | Upregulated |
| Rat | fpgs | 1.550359 | Upregulated |
| Rat | ubr1 | 1.549271 | Upregulated |
| Rat | kcnj2 | 1.529666 | Upregulated |
| Rat | ntn1 | 1.500451 | Upregulated |
| Rat | rhox13 | 1.500434 | Upregulated |
| Rat | krt24 | 1.481836 | Upregulated |
| Rat | btla | 1.475586 | Upregulated |
| Rat | tmc5 | 1.469893 | Upregulated |
| Rat | acsm4 | 1.464534 | Upregulated |
| Rat | il5 | 1.460111 | Upregulated |
| Rat | cxcl9 | 1.457707 | Upregulated |
| Rat | gstt4 | 1.439882 | Upregulated |
| Rat | crybb2 | 1.424533 | Upregulated |
| Rat | or1e16 | 1.420792 | Upregulated |
| Rat | gpx5 | 1.420031 | Upregulated |
| Rat | tns4 | 1.412455 | Upregulated |
| Rat | slc26a5 | 1.407663 | Upregulated |
| Rat | atp2c2 | 1.40579 | Upregulated |
| Rat | adgrf3 | 1.404591 | Upregulated |
| Rat | npffr2 | 1.401888 | Upregulated |
| Rat | ubxn10 | 1.39885 | Upregulated |
| Mouse | hspa1a | -3.23 | Downregulated |
| Mouse | ccl3 | -2.57019 | Downregulated |
| Mouse | fosb | -2.18223 | Downregulated |
| Mouse | npas4 | -2 | Downregulated |
| Mouse | cxcl1 | -1.93932 | Downregulated |
| Mouse | cd14 | -1.83 | Downregulated |
| Mouse | fos | -1.8 | Downregulated |
| Mouse | atf3 | -1.69217 | Downregulated |
| Mouse | ccn1 | -1.605 | Downregulated |
| Mouse | ptgs2 | -1.45069 | Downregulated |
| Mouse | hmox1 | -1.37 | Downregulated |
| Mouse | gadd45g | -1.36956 | Downregulated |
| Mouse | gadd45b | -1.3656 | Downregulated |
| Mouse | zfp36 | -1.35376 | Downregulated |
| Mouse | jun | -1.33962 | Downregulated |
| Mouse | apold1 | -1.27103 | Downregulated |
| Mouse | socs3 | -1.2665 | Downregulated |
| Mouse | gpr84 | -1.244 | Downregulated |
| Mouse | cdkn1a | -1.2393 | Downregulated |
| Mouse | ccl12 | -1.20469 | Downregulated |
| Mouse | ccl2 | -1.19933 | Downregulated |
| Mouse | hspb1 | -1.10979 | Downregulated |
| Mouse | adamts1 | -1.1 | Downregulated |
| Mouse | slc15a3 | -0.97717 | Downregulated |
| Mouse | nptx2 | -0.97553 | Downregulated |
| Mouse | osmr | -0.96819 | Downregulated |
| Mouse | ccl9 | -0.96085 | Downregulated |
| Mouse | ctla2a | -0.92368 | Downregulated |
| Mouse | cxcl10 | -0.90992 | Downregulated |
| Mouse | baz1a | -0.90483 | Downregulated |
| Rat | cxcl2 | -5.79947 | Downregulated |
| Rat | hspa1a | -5.56245 | Downregulated |
| Rat | cxcl1 | -5.28367 | Downregulated |
| Rat | ccl3 | -4.13 | Downregulated |
| Rat | npas4 | -3.91 | Downregulated |
| Rat | fosl1 | -3.12707 | Downregulated |
| Rat | fos | -3.05638 | Downregulated |
| Rat | gdf15 | -3.05428 | Downregulated |
| Rat | ccl2 | -2.64 | Downregulated |
| Rat | tfpi2 | -2.51766 | Downregulated |
| Rat | ccn1 | -2.38719 | Downregulated |
| Rat | olr1 | -2.38 | Downregulated |
| Rat | arc | -2.11093 | Downregulated |
| Rat | ptges | -2.06713 | Downregulated |
| Rat | gadd45g | -2.06554 | Downregulated |
| Rat | ccl4 | -2.02087 | Downregulated |
| Rat | sele | -1.96454 | Downregulated |
| Rat | btg2 | -1.88634 | Downregulated |
| Rat | atf3 | -1.8 | Downregulated |
| Rat | cxcl10 | -1.73596 | Downregulated |
| Rat | zfp36 | -1.72 | Downregulated |
| Rat | fosb | -1.66462 | Downregulated |
| Rat | spp1 | -1.64059 | Downregulated |
| Rat | dusp5 | -1.62718 | Downregulated |
| Rat | apold1 | -1.61835 | Downregulated |
| Rat | serpina3n | -1.60606 | Downregulated |
| Rat | cebpd | -1.54399 | Downregulated |
| Rat | nr4a1 | -1.52 | Downregulated |
| Rat | timp1 | -1.49433 | Downregulated |
| Rat | egr2 | -1.47085 | Downregulated |

**Suppl. Table 3:** Median fold changes of most differentially expressed genes subacutely after stroke in mice and rats

| **Species** | **Gene.symbol** | **logFC** | **Regulation** |
| --- | --- | --- | --- |
| Mouse | or5p81 | 2.7 | Upregulated |
| Mouse | or51e1 | 2.47 | Upregulated |
| Mouse | iho1 | 2.45 | Upregulated |
| Mouse | or51r1 | 2.43 | Upregulated |
| Mouse | lbhd2 | 2.32 | Upregulated |
| Mouse | ccn3 | 2.25 | Upregulated |
| Mouse | gpr88 | 2.17 | Upregulated |
| Mouse | ripor2 | 1.99 | Upregulated |
| Mouse | vxn | 1.88 | Upregulated |
| Mouse | pnma8b | 1.77 | Upregulated |
| Mouse | septin5 | 1.75 | Upregulated |
| Mouse | ankrd63 | 1.698484 | Upregulated |
| Mouse | samd13 | 1.69 | Upregulated |
| Mouse | prr33 | 1.68805 | Upregulated |
| Mouse | armh4 | 1.66 | Upregulated |
| Mouse | or6b6 | 1.65 | Upregulated |
| Mouse | lrrc10b | 1.601248 | Upregulated |
| Mouse | penk | 1.600141 | Upregulated |
| Mouse | or51f2 | 1.6 | Upregulated |
| Mouse | pmch | 1.584317 | Upregulated |
| Mouse | minar2 | 1.55 | Upregulated |
| Mouse | pnma8a | 1.51 | Upregulated |
| Mouse | carmil3 | 1.5 | Upregulated |
| Mouse | tmem35a | 1.5 | Upregulated |
| Mouse | myorg | 1.48 | Upregulated |
| Mouse | gpr101 | 1.473394 | Upregulated |
| Mouse | il36rn | 1.44 | Upregulated |
| Mouse | agt | 1.4 | Upregulated |
| Mouse | magee2 | 1.3931 | Upregulated |
| Mouse | cfap20dc | 1.38 | Upregulated |
| Rat | llcfc1 | 1.543384 | Upregulated |
| Rat | or4c29 | 1.444568 | Upregulated |
| Rat | spem2 | 1.429851 | Upregulated |
| Rat | or10s1 | 1.427641 | Upregulated |
| Rat | or4c1 | 1.329588 | Upregulated |
| Rat | plscr5 | 1.321666 | Upregulated |
| Rat | drd2 | 1.291753 | Upregulated |
| Rat | syndig1l | 1.223672 | Upregulated |
| Rat | ccdc169 | 1.164295 | Upregulated |
| Rat | serpinb13 | 1.149096 | Upregulated |
| Rat | slc35d3 | 1.141844 | Upregulated |
| Rat | or9m1b | 1.100771 | Upregulated |
| Rat | gpr88 | 1.08 | Upregulated |
| Rat | ccn3 | 1.057931 | Upregulated |
| Rat | tmem269 | 1.0541 | Upregulated |
| Rat | or2m12 | 1.021211 | Upregulated |
| Rat | or2y16 | 1.019092 | Upregulated |
| Rat | tas2r104 | 1.012584 | Upregulated |
| Rat | atp13a4 | 1.00878 | Upregulated |
| Rat | fezf1 | 0.98549 | Upregulated |
| Rat | or5w13 | 0.984987 | Upregulated |
| Rat | or4c110 | 0.979133 | Upregulated |
| Rat | or8s8 | 0.97569 | Upregulated |
| Rat | tmem151b | 0.970617 | Upregulated |
| Rat | myorg | 0.968009 | Upregulated |
| Rat | lamp5 | 0.967186 | Upregulated |
| Rat | septin5 | 0.952285 | Upregulated |
| Rat | kics2 | 0.948113 | Upregulated |
| Rat | drd1 | 0.943372 | Upregulated |
| Rat | slc15a5 | 0.940421 | Upregulated |
| Mouse | il36g | -5.25 | Downregulated |
| Mouse | lcn2 | -4.67 | Downregulated |
| Mouse | spp1 | -4.34252 | Downregulated |
| Mouse | hspa1a | -3.40933 | Downregulated |
| Mouse | lgals3 | -3.37851 | Downregulated |
| Mouse | or5m13b | -3.29 | Downregulated |
| Mouse | timp1 | -3.26356 | Downregulated |
| Mouse | hmox1 | -3.25 | Downregulated |
| Mouse | ccl3 | -3.06095 | Downregulated |
| Mouse | ccl12 | -3.05225 | Downregulated |
| Mouse | atf3 | -2.96352 | Downregulated |
| Mouse | gsta5 | -2.945 | Downregulated |
| Mouse | mmp3 | -2.93996 | Downregulated |
| Mouse | garin4 | -2.89 | Downregulated |
| Mouse | il6 | -2.79909 | Downregulated |
| Mouse | ccn1 | -2.75 | Downregulated |
| Mouse | ccl4 | -2.7288 | Downregulated |
| Mouse | cep295nl | -2.58 | Downregulated |
| Mouse | il11 | -2.55229 | Downregulated |
| Mouse | s100a11 | -2.54592 | Downregulated |
| Mouse | or9q1 | -2.54 | Downregulated |
| Mouse | s100a8 | -2.50474 | Downregulated |
| Mouse | tgm1 | -2.49 | Downregulated |
| Mouse | s100a9 | -2.40648 | Downregulated |
| Mouse | tgfbi | -2.38389 | Downregulated |
| Mouse | ccl9 | -2.37 | Downregulated |
| Mouse | vim | -2.36415 | Downregulated |
| Mouse | socs3 | -2.3613 | Downregulated |
| Mouse | cd14 | -2.35361 | Downregulated |
| Mouse | arpc1b | -2.33659 | Downregulated |
| Rat | ccl2 | -4.71 | Downregulated |
| Rat | msr1 | -3.93519 | Downregulated |
| Rat | spp1 | -3.83 | Downregulated |
| Rat | lgals3 | -3.69 | Downregulated |
| Rat | cd300c2 | -3.585 | Downregulated |
| Rat | timp1 | -3.48437 | Downregulated |
| Rat | hspb1 | -3.38578 | Downregulated |
| Rat | rrm2 | -3.35 | Downregulated |
| Rat | gpnmb | -3.18335 | Downregulated |
| Rat | clec7a | -2.915 | Downregulated |
| Rat | pbk | -2.77482 | Downregulated |
| Rat | lcn2 | -2.76283 | Downregulated |
| Rat | hspa1a | -2.67 | Downregulated |
| Rat | cd14 | -2.63671 | Downregulated |
| Rat | ndc80 | -2.52468 | Downregulated |
| Rat | pimreg | -2.50052 | Downregulated |
| Rat | ckap2 | -2.49427 | Downregulated |
| Rat | gpr183 | -2.41226 | Downregulated |
| Rat | hmox1 | -2.39495 | Downregulated |
| Rat | ccl7 | -2.38 | Downregulated |
| Rat | serpina3n | -2.375 | Downregulated |
| Rat | slamf9 | -2.36 | Downregulated |
| Rat | siglec1 | -2.35 | Downregulated |
| Rat | plp2 | -2.34472 | Downregulated |
| Rat | cdk1 | -2.34 | Downregulated |
| Rat | prc1 | -2.32972 | Downregulated |
| Rat | serpine1 | -2.3275 | Downregulated |
| Rat | hk3 | -2.31937 | Downregulated |
| Rat | glipr2 | -2.27281 | Downregulated |
| Rat | crisp3 | -2.25925 | Downregulated |

**Suppl. Table 4:** Median fold changes of most differentially expressed genes subacutely after stroke in mice and rats

| **Species** | **Gene.symbol** | **logFC** | **Regulation** |
| --- | --- | --- | --- |
| Mouse | ttr | 4.604395 | Upregulated |
| Mouse | slc4a5 | 3.074951 | Upregulated |
| Mouse | tmem72 | 3.057662 | Upregulated |
| Mouse | f5 | 2.85722 | Upregulated |
| Mouse | shox2 | 2.742124 | Upregulated |
| Mouse | kcne2 | 2.563359 | Upregulated |
| Mouse | tph2 | 2.369253 | Upregulated |
| Mouse | otx2 | 2.219709 | Upregulated |
| Mouse | cldn2 | 1.920661 | Upregulated |
| Mouse | prlr | 1.832764 | Upregulated |
| Mouse | slc18a2 | 1.681306 | Upregulated |
| Mouse | folr1 | 1.676182 | Upregulated |
| Mouse | glra1 | 1.671926 | Upregulated |
| Mouse | clic6 | 1.639709 | Upregulated |
| Mouse | sostdc1 | 1.595481 | Upregulated |
| Mouse | kl | 1.504896 | Upregulated |
| Mouse | heatr5b | 1.397912 | Upregulated |
| Mouse | hic2 | 1.32883 | Upregulated |
| Mouse | kdr | 1.305641 | Upregulated |
| Mouse | egr2 | 1.298065 | Upregulated |
| Mouse | colq | 1.297019 | Upregulated |
| Mouse | chrnb3 | 1.283019 | Upregulated |
| Mouse | kcnj13 | 1.267239 | Upregulated |
| Mouse | cyyr1 | 1.225586 | Upregulated |
| Mouse | ebf3 | 1.197745 | Upregulated |
| Mouse | npas4 | 1.184569 | Upregulated |
| Mouse | slc6a5 | 1.180536 | Upregulated |
| Mouse | irx3 | 1.15653 | Upregulated |
| Mouse | fos | 1.139812 | Upregulated |
| Mouse | dgkk | 1.132489 | Upregulated |
| Rat | msr1 | 5.189444 | Upregulated |
| Rat | cd300c2 | 5.144992 | Upregulated |
| Rat | stra6l | 4.002048 | Upregulated |
| Rat | ckap2l | 3.532761 | Upregulated |
| Rat | pclaf | 3.522151 | Upregulated |
| Rat | aoah | 3.143599 | Upregulated |
| Rat | rexo5 | 3.052205 | Upregulated |
| Rat | ticam2 | 2.703448 | Upregulated |
| Rat | cd84 | 2.52246 | Upregulated |
| Rat | pimreg | 2.486643 | Upregulated |
| Rat | nlrp3 | 2.435346 | Upregulated |
| Rat | lrmda | 2.39649 | Upregulated |
| Rat | ugt1a2 | 2.339259 | Upregulated |
| Rat | tmem273 | 2.224012 | Upregulated |
| Rat | tent5c | 2.102321 | Upregulated |
| Rat | mastl | 2.074469 | Upregulated |
| Rat | ckap2 | 2.01047 | Upregulated |
| Rat | exph5 | 2.00942 | Upregulated |
| Rat | dbf4 | 1.970916 | Upregulated |
| Rat | chrna3 | 1.94506 | Upregulated |
| Rat | lamp5 | 1.934505 | Upregulated |
| Rat | ap3b2 | 1.89332 | Upregulated |
| Rat | arc | 1.86416 | Upregulated |
| Rat | pheta2 | 1.835711 | Upregulated |
| Rat | pla2g4e | 1.74834 | Upregulated |
| Rat | dock3 | 1.69326 | Upregulated |
| Rat | car10 | 1.67326 | Upregulated |
| Rat | tcim | 1.669018 | Upregulated |
| Rat | fam19a1 | 1.62812 | Upregulated |
| Rat | tph1 | 1.6 | Upregulated |
| Mouse | lcn2 | -5.86277 | Downregulated |
| Mouse | mmp12 | -4.94706 | Downregulated |
| Mouse | mmp13 | -4.71202 | Downregulated |
| Mouse | clec7a | -4.6029 | Downregulated |
| Mouse | lyz2 | -4.50419 | Downregulated |
| Mouse | gpnmb | -4.21028 | Downregulated |
| Mouse | lgals3 | -4.0741 | Downregulated |
| Mouse | atp6v0d2 | -4.05053 | Downregulated |
| Mouse | itgax | -3.90269 | Downregulated |
| Mouse | cd5l | -3.79819 | Downregulated |
| Mouse | cst7 | -3.6978 | Downregulated |
| Mouse | ms4a15 | -3.63077 | Downregulated |
| Mouse | steap4 | -3.61647 | Downregulated |
| Mouse | tgm1 | -3.57605 | Downregulated |
| Mouse | spp1 | -3.32137 | Downregulated |
| Mouse | cd74 | -3.2289 | Downregulated |
| Mouse | mmp3 | -3.15049 | Downregulated |
| Mouse | a2m | -3.14265 | Downregulated |
| Mouse | apoc1 | -3.09946 | Downregulated |
| Mouse | kcnj15 | -3.03341 | Downregulated |
| Mouse | eomes | -2.8596 | Downregulated |
| Mouse | lum | -2.82412 | Downregulated |
| Mouse | cybb | -2.81514 | Downregulated |
| Mouse | col1a1 | -2.76947 | Downregulated |
| Mouse | c3 | -2.72523 | Downregulated |
| Mouse | cd22 | -2.69395 | Downregulated |
| Mouse | vipr2 | -2.62996 | Downregulated |
| Mouse | tbx21 | -2.61219 | Downregulated |
| Mouse | timp1 | -2.5424 | Downregulated |
| Mouse | cdhr1 | -2.49341 | Downregulated |
| Rat | clec7a | -5.44062 | Downregulated |
| Rat | tlr8 | -4.6066 | Downregulated |
| Rat | itgax | -4.59114 | Downregulated |
| Rat | evi2b | -4.55916 | Downregulated |
| Rat | fam46c | -4.40754 | Downregulated |
| Rat | cenpe | -4.33966 | Downregulated |
| Rat | mki67 | -4.27138 | Downregulated |
| Rat | wdfy4 | -4.12 | Downregulated |
| Rat | ddx60 | -3.75954 | Downregulated |
| Rat | ccl6 | -3.72159 | Downregulated |
| Rat | igkc | -3.6946 | Downregulated |
| Rat | spp1 | -3.67 | Downregulated |
| Rat | gpnmb | -3.62183 | Downregulated |
| Rat | itih2 | -3.60034 | Downregulated |
| Rat | card14 | -3.52862 | Downregulated |
| Rat | lum | -3.49 | Downregulated |
| Rat | irf8 | -3.48909 | Downregulated |
| Rat | frrs1 | -3.47068 | Downregulated |
| Rat | clec4a3 | -3.45617 | Downregulated |
| Rat | cd180 | -3.43595 | Downregulated |
| Rat | cd33 | -3.419 | Downregulated |
| Rat | mcub | -3.12736 | Downregulated |
| Rat | cnn2 | -3.07354 | Downregulated |
| Rat | golm1 | -3.05617 | Downregulated |
| Rat | shcbp1 | -3.04251 | Downregulated |
| Rat | col4a6 | -3.00379 | Downregulated |
| Rat | tmem173 | -2.93732 | Downregulated |
| Rat | col8a2 | -2.93364 | Downregulated |
| Rat | fam105a | -2.92792 | Downregulated |
| Rat | lgals3 | -2.91928 | Downregulated |

**Suppl. Table 5**: Median fold changes of 60 most differentially expressed genes acutely following tMCAo and pMCAo

| **Stroke.model** | **Gene.symbol** | **logFC** | **Regulation** |
| --- | --- | --- | --- |
| pMCAO | or1e16 | 1.414737 | Upregulated |
| pMCAO | krt9 | 1.22536 | Upregulated |
| pMCAO | glra3 | 1.183348 | Upregulated |
| pMCAO | crnn | 1.178879 | Upregulated |
| pMCAO | gpr33 | 1.176362 | Upregulated |
| pMCAO | fcrl6 | 1.156481 | Upregulated |
| pMCAO | cfap276 | 1.151251 | Upregulated |
| pMCAO | chrm5 | 1.133946 | Upregulated |
| pMCAO | ccdc88c | 1.127058 | Upregulated |
| pMCAO | krt82 | 1.10973 | Upregulated |
| pMCAO | dnah9 | 1.086953 | Upregulated |
| pMCAO | or12d17 | 1.085129 | Upregulated |
| pMCAO | gnrhr | 1.084862 | Upregulated |
| pMCAO | rhox12 | 1.069734 | Upregulated |
| pMCAO | obp2b | 1.069565 | Upregulated |
| pMCAO | prop1 | 1.05079 | Upregulated |
| pMCAO | calm5 | 1.04 | Upregulated |
| pMCAO | a1cf | 0.988236 | Upregulated |
| pMCAO | cyct | 0.968922 | Upregulated |
| pMCAO | neurog1 | 0.961225 | Upregulated |
| pMCAO | sbk3 | 0.957331 | Upregulated |
| pMCAO | gm1979 | 0.954 | Upregulated |
| pMCAO | slc26a5 | 0.946118 | Upregulated |
| pMCAO | amy2a3 | 0.94589 | Upregulated |
| pMCAO | olah | 0.927742 | Upregulated |
| pMCAO | lctl | 0.926399 | Upregulated |
| pMCAO | atp1a4 | 0.919963 | Upregulated |
| pMCAO | smagp | 0.91101 | Upregulated |
| pMCAO | efcab3 | 0.910366 | Upregulated |
| pMCAO | klra22 | 0.908047 | Upregulated |
| tMCAO | gm20852 | 1.501318 | Upregulated |
| tMCAO | gm20867 | 1.491161 | Upregulated |
| tMCAO | gm20865 | 1.39704 | Upregulated |
| tMCAO | or51q1c | 1.387311 | Upregulated |
| tMCAO | or5w19 | 1.327785 | Upregulated |
| tMCAO | prl2c2 | 1.263611 | Upregulated |
| tMCAO | pramel50 | 1.240501 | Upregulated |
| tMCAO | gm16405 | 1.20659 | Upregulated |
| tMCAO | or6c6 | 1.205907 | Upregulated |
| tMCAO | gm20871 | 1.179168 | Upregulated |
| tMCAO | gm13275 | 1.150681 | Upregulated |
| tMCAO | gm20822 | 1.146521 | Upregulated |
| tMCAO | gm10488 | 1.143591 | Upregulated |
| tMCAO | gm20809 | 1.139693 | Upregulated |
| tMCAO | or6d15 | 1.129099 | Upregulated |
| tMCAO | or51q1 | 1.112997 | Upregulated |
| tMCAO | vmn1r104 | 1.102398 | Upregulated |
| tMCAO | or14j2 | 1.097793 | Upregulated |
| tMCAO | sprr2a2 | 1.097743 | Upregulated |
| tMCAO | gm13287 | 1.094107 | Upregulated |
| tMCAO | or1j13 | 1.077981 | Upregulated |
| tMCAO | gm6367 | 1.065063 | Upregulated |
| tMCAO | gm10406 | 1.05209 | Upregulated |
| tMCAO | btbd35f12 | 1.050113 | Upregulated |
| tMCAO | vmn1r131 | 1.049586 | Upregulated |
| tMCAO | or5p80 | 1.039123 | Upregulated |
| tMCAO | slxl1 | 1.024773 | Upregulated |
| tMCAO | or9g19 | 1.02319 | Upregulated |
| tMCAO | gm13277 | 1.022258 | Upregulated |
| tMCAO | gm13290 | 1.022258 | Upregulated |
| pMCAO | cxcl1 | -5.16342 | Downregulated |
| pMCAO | hspa1b | -4.21 | Downregulated |
| pMCAO | hspa1a | -3.675 | Downregulated |
| pMCAO | npas4 | -3.395 | Downregulated |
| pMCAO | ccl3 | -3.175 | Downregulated |
| pMCAO | fos | -2.5656 | Downregulated |
| pMCAO | ccl2 | -2.24701 | Downregulated |
| pMCAO | fosb | -1.96356 | Downregulated |
| pMCAO | lilrb4a | -1.935 | Downregulated |
| pMCAO | gadd45g | -1.77471 | Downregulated |
| pMCAO | ccn1 | -1.64 | Downregulated |
| pMCAO | cd14 | -1.59108 | Downregulated |
| pMCAO | gadd45b | -1.55827 | Downregulated |
| pMCAO | ccl4 | -1.53103 | Downregulated |
| pMCAO | inhba | -1.52152 | Downregulated |
| pMCAO | atf3 | -1.49202 | Downregulated |
| pMCAO | apold1 | -1.48418 | Downregulated |
| pMCAO | dusp5 | -1.41742 | Downregulated |
| pMCAO | nptx2 | -1.34336 | Downregulated |
| pMCAO | hspb1 | -1.29 | Downregulated |
| pMCAO | jun | -1.20909 | Downregulated |
| pMCAO | timp1 | -1.17795 | Downregulated |
| pMCAO | ptgs2 | -1.17529 | Downregulated |
| pMCAO | il1a | -1.16556 | Downregulated |
| pMCAO | egr2 | -1.11837 | Downregulated |
| pMCAO | gdf15 | -1.11835 | Downregulated |
| pMCAO | junb | -1.09138 | Downregulated |
| pMCAO | btg2 | -1.07 | Downregulated |
| pMCAO | nfil3 | -1.03956 | Downregulated |
| pMCAO | arc | -1.02 | Downregulated |
| tMCAO | ccn1 | -3.2998 | Downregulated |
| tMCAO | hspa1b | -3.15162 | Downregulated |
| tMCAO | ccl3 | -2.93092 | Downregulated |
| tMCAO | hspa1a | -2.60051 | Downregulated |
| tMCAO | fos | -2.15352 | Downregulated |
| tMCAO | cd14 | -2.12598 | Downregulated |
| tMCAO | ccl4 | -2.05179 | Downregulated |
| tMCAO | lcn2 | -1.9626 | Downregulated |
| tMCAO | atf3 | -1.95051 | Downregulated |
| tMCAO | npas4 | -1.85169 | Downregulated |
| tMCAO | cxcl1 | -1.79691 | Downregulated |
| tMCAO | adamts1 | -1.75963 | Downregulated |
| tMCAO | timp1 | -1.75324 | Downregulated |
| tMCAO | hmox1 | -1.68282 | Downregulated |
| tMCAO | hspb1 | -1.65899 | Downregulated |
| tMCAO | ccl2 | -1.6467 | Downregulated |
| tMCAO | socs3 | -1.64074 | Downregulated |
| tMCAO | apold1 | -1.62406 | Downregulated |
| tMCAO | zfp36 | -1.55668 | Downregulated |
| tMCAO | ccl9 | -1.55476 | Downregulated |
| tMCAO | cxcl10 | -1.41101 | Downregulated |
| tMCAO | gadd45g | -1.39167 | Downregulated |
| tMCAO | ch25h | -1.39044 | Downregulated |
| tMCAO | ccl12 | -1.37854 | Downregulated |
| tMCAO | gpr84 | -1.36506 | Downregulated |
| tMCAO | ier2 | -1.34434 | Downregulated |
| tMCAO | lgals3 | -1.31987 | Downregulated |
| tMCAO | depp1 | -1.31042 | Downregulated |
| tMCAO | fosb | -1.31024 | Downregulated |
| tMCAO | egr2 | -1.29178 | Downregulated |

**Suppl. Table 6:** Median fold changes of 60 most differentially expressed genes following tMCAo and pMCAo and photothrombotic stroke (PT) at the long-term period.

| **Stroke.model** | **Gene.symbol** | **logFC** | **Regulation** |
| --- | --- | --- | --- |
| PT | ttr | 4.604395 | Upregulated |
| PT | slc4a5 | 3.074951 | Upregulated |
| PT | tmem72 | 3.057662 | Upregulated |
| PT | f5 | 2.85722 | Upregulated |
| PT | shox2 | 2.742124 | Upregulated |
| PT | kcne2 | 2.563359 | Upregulated |
| PT | tph2 | 2.369253 | Upregulated |
| PT | otx2 | 2.219709 | Upregulated |
| PT | cldn2 | 1.920661 | Upregulated |
| PT | prlr | 1.832764 | Upregulated |
| PT | slc18a2 | 1.681306 | Upregulated |
| PT | folr1 | 1.676182 | Upregulated |
| PT | glra1 | 1.671926 | Upregulated |
| PT | clic6 | 1.639709 | Upregulated |
| PT | sostdc1 | 1.595481 | Upregulated |
| PT | kl | 1.504896 | Upregulated |
| PT | kdr | 1.305641 | Upregulated |
| PT | egr2 | 1.298065 | Upregulated |
| PT | colq | 1.297019 | Upregulated |
| PT | chrnb3 | 1.283019 | Upregulated |
| PT | kcnj13 | 1.267239 | Upregulated |
| PT | cyyr1 | 1.225586 | Upregulated |
| PT | ebf3 | 1.197745 | Upregulated |
| PT | npas4 | 1.184569 | Upregulated |
| PT | slc6a5 | 1.180536 | Upregulated |
| PT | irx3 | 1.15653 | Upregulated |
| PT | fos | 1.139812 | Upregulated |
| PT | dgkk | 1.132489 | Upregulated |
| PT | flt1 | 1.1077 | Upregulated |
| PT | dll4 | 1.09989 | Upregulated |
| pMCAO | spp1 | 8.264112 | Upregulated |
| pMCAO | mmp7 | 6.2752 | Upregulated |
| pMCAO | cd8a | 5.982131 | Upregulated |
| pMCAO | ccl2 | 5.924804 | Upregulated |
| pMCAO | fabp4 | 5.88733 | Upregulated |
| pMCAO | ccl7 | 5.865053 | Upregulated |
| pMCAO | fcnb | 5.830286 | Upregulated |
| pMCAO | lgals3 | 5.705464 | Upregulated |
| pMCAO | hk3 | 5.495872 | Upregulated |
| pMCAO | gpnmb | 5.476778 | Upregulated |
| pMCAO | mmp12 | 5.39491 | Upregulated |
| pMCAO | slpi | 5.376632 | Upregulated |
| pMCAO | plau | 5.349219 | Upregulated |
| pMCAO | slamf9 | 5.343721 | Upregulated |
| pMCAO | lilrb4 | 5.256452 | Upregulated |
| pMCAO | knstrn | 5.245167 | Upregulated |
| pMCAO | msr1 | 5.189444 | Upregulated |
| pMCAO | cd300c2 | 5.144992 | Upregulated |
| pMCAO | loc689770 | 5.116217 | Upregulated |
| pMCAO | cdk1 | 5.06401 | Upregulated |
| pMCAO | cenpa | 5.021671 | Upregulated |
| pMCAO | lcn2 | 4.969105 | Upregulated |
| pMCAO | kif2c | 4.929121 | Upregulated |
| pMCAO | cd68 | 4.824014 | Upregulated |
| pMCAO | c9h6orf141 | 4.742209 | Upregulated |
| pMCAO | igsf6 | 4.672426 | Upregulated |
| pMCAO | pbk | 4.672118 | Upregulated |
| pMCAO | cdca2 | 4.664077 | Upregulated |
| pMCAO | prc1 | 4.630398 | Upregulated |
| pMCAO | il1rl1 | 4.600377 | Upregulated |
| tMCAO | arc | 3.01632 | Upregulated |
| tMCAO | ccn1 | 2.393321 | Upregulated |
| tMCAO | exph5 | 2.00942 | Upregulated |
| tMCAO | nr4a1 | 1.99068 | Upregulated |
| tMCAO | lamp5 | 1.961162 | Upregulated |
| tMCAO | hbb-b1 | 1.952817 | Upregulated |
| tMCAO | chrna3 | 1.94506 | Upregulated |
| tMCAO | kcnv1 | 1.833551 | Upregulated |
| tMCAO | cort | 1.774546 | Upregulated |
| tMCAO | pla2g4e | 1.74834 | Upregulated |
| tMCAO | crh | 1.729589 | Upregulated |
| tMCAO | hbb | 1.70418 | Upregulated |
| tMCAO | egr2 | 1.68452 | Upregulated |
| tMCAO | lrriq1 | 1.629356 | Upregulated |
| tMCAO | egr4 | 1.62226 | Upregulated |
| tMCAO | chrm1 | 1.6185 | Upregulated |
| tMCAO | tph1 | 1.6 | Upregulated |
| tMCAO | sowahb | 1.56956 | Upregulated |
| tMCAO | sept5 | 1.56708 | Upregulated |
| tMCAO | vom2r40 | 1.5605 | Upregulated |
| tMCAO | cplx2 | 1.46798 | Upregulated |
| tMCAO | wnt10a | 1.452479 | Upregulated |
| tMCAO | sptssb | 1.449895 | Upregulated |
| tMCAO | tac1 | 1.44 | Upregulated |
| tMCAO | nefm | 1.42614 | Upregulated |
| tMCAO | fam43b | 1.41272 | Upregulated |
| tMCAO | egr1 | 1.393349 | Upregulated |
| tMCAO | hes2 | 1.37405 | Upregulated |
| tMCAO | hbb-bs | 1.366359 | Upregulated |
| tMCAO | mslnl | 1.35796 | Upregulated |
| PT | lcn2 | -5.86277 | Downregulated |
| PT | c3 | -5.52345 | Downregulated |
| PT | mmp12 | -4.94706 | Downregulated |
| PT | mmp13 | -4.71202 | Downregulated |
| PT | clec7a | -4.6029 | Downregulated |
| PT | lyz2 | -4.50419 | Downregulated |
| PT | gpnmb | -4.21028 | Downregulated |
| PT | lgals3 | -4.0741 | Downregulated |
| PT | atp6v0d2 | -4.05053 | Downregulated |
| PT | itgax | -3.90269 | Downregulated |
| PT | cd5l | -3.79819 | Downregulated |
| PT | cst7 | -3.6978 | Downregulated |
| PT | ms4a15 | -3.63077 | Downregulated |
| PT | steap4 | -3.61647 | Downregulated |
| PT | col3a1 | -3.59967 | Downregulated |
| PT | tgm1 | -3.57605 | Downregulated |
| PT | spp1 | -3.32137 | Downregulated |
| PT | cd74 | -3.2289 | Downregulated |
| PT | mmp3 | -3.15049 | Downregulated |
| PT | a2m | -3.14265 | Downregulated |
| PT | apoc1 | -3.09946 | Downregulated |
| PT | kcnj15 | -3.03341 | Downregulated |
| PT | eomes | -2.8596 | Downregulated |
| PT | lum | -2.82412 | Downregulated |
| PT | cybb | -2.81514 | Downregulated |
| PT | col1a1 | -2.76947 | Downregulated |
| PT | cd22 | -2.69395 | Downregulated |
| PT | vipr2 | -2.62996 | Downregulated |
| PT | tbx21 | -2.61219 | Downregulated |
| PT | timp1 | -2.5424 | Downregulated |
| pMCAO | arc | -2.4609 | Downregulated |
| pMCAO | tnnc2 | -2.40596 | Downregulated |
| pMCAO | slc12a1 | -2.35463 | Downregulated |
| pMCAO | nr4a1 | -2.21391 | Downregulated |
| pMCAO | tob1 | -2.17245 | Downregulated |
| pMCAO | selenov | -2.08931 | Downregulated |
| pMCAO | ckmt2 | -2.03497 | Downregulated |
| pMCAO | arr3 | -2.03236 | Downregulated |
| pMCAO | egr4 | -1.9534 | Downregulated |
| pMCAO | lamp5 | -1.89364 | Downregulated |
| pMCAO | nr4a3 | -1.88436 | Downregulated |
| pMCAO | cftr | -1.80832 | Downregulated |
| pMCAO | pcdh20 | -1.74008 | Downregulated |
| pMCAO | dusp1 | -1.72264 | Downregulated |
| pMCAO | alb | -1.71949 | Downregulated |
| pMCAO | nphp4 | -1.67489 | Downregulated |
| pMCAO | cbln4 | -1.66329 | Downregulated |
| pMCAO | cbln2 | -1.64809 | Downregulated |
| pMCAO | slc17a6 | -1.64578 | Downregulated |
| pMCAO | or10g9b | -1.62866 | Downregulated |
| pMCAO | egr2 | -1.61769 | Downregulated |
| pMCAO | rnd1 | -1.61187 | Downregulated |
| pMCAO | cartpt | -1.60549 | Downregulated |
| pMCAO | vsnl1 | -1.59117 | Downregulated |
| pMCAO | crh | -1.57901 | Downregulated |
| pMCAO | khdrbs2 | -1.57816 | Downregulated |
| pMCAO | qrfpr | -1.57744 | Downregulated |
| pMCAO | mrgprd | -1.56501 | Downregulated |
| pMCAO | kcnj3 | -1.55445 | Downregulated |
| pMCAO | pnlip | -1.55204 | Downregulated |
| tMCAO | spp1 | -6.70702 | Downregulated |
| tMCAO | clec7a | -5.44062 | Downregulated |
| tMCAO | lilrb4 | -5.01709 | Downregulated |
| tMCAO | tlr8 | -4.6066 | Downregulated |
| tMCAO | itgax | -4.59114 | Downregulated |
| tMCAO | evi2b | -4.55916 | Downregulated |
| tMCAO | mmp12 | -4.423 | Downregulated |
| tMCAO | cenpe | -4.33966 | Downregulated |
| tMCAO | mki67 | -4.27138 | Downregulated |
| tMCAO | lum | -4.2251 | Downregulated |
| tMCAO | cd68 | -4.14 | Downregulated |
| tMCAO | wdfy4 | -4.12 | Downregulated |
| tMCAO | gpnmb | -3.99 | Downregulated |
| tMCAO | fcrl2 | -3.7672 | Downregulated |
| tMCAO | ccl6 | -3.72159 | Downregulated |
| tMCAO | igkc | -3.6946 | Downregulated |
| tMCAO | ifitm1 | -3.69 | Downregulated |
| tMCAO | itih2 | -3.60034 | Downregulated |
| tMCAO | fabp4 | -3.56505 | Downregulated |
| tMCAO | arl11 | -3.56278 | Downregulated |
| tMCAO | card14 | -3.52862 | Downregulated |
| tMCAO | irf8 | -3.48909 | Downregulated |
| tMCAO | frrs1 | -3.47068 | Downregulated |
| tMCAO | clec4a3 | -3.45617 | Downregulated |
| tMCAO | cd180 | -3.43595 | Downregulated |
| tMCAO | lgals3 | -3.42 | Downregulated |
| tMCAO | cd33 | -3.419 | Downregulated |
| tMCAO | blnk | -3.30336 | Downregulated |
| tMCAO | bcl2a1 | -3.24935 | Downregulated |
| tMCAO | plau | -3.22862 | Downregulated |

**Suppl. Table 7:** Median fold changes of 60 most differentially expressed genes at different time periods after stroke.

| **Time.points** | **Gene.symbol** | **logFC** | **Regulation** |
| --- | --- | --- | --- |
| acute | mir374b | 2.15 | Upregulated |
| acute | mir421 | 2.1 | Upregulated |
| acute | mir543 | 1.803585 | Upregulated |
| acute | mir329 | 1.76 | Upregulated |
| acute | cxcl6 | 1.620217 | Upregulated |
| acute | rgd1564571 | 1.533229 | Upregulated |
| acute | mgc114246 | 1.483475 | Upregulated |
| acute | loc286992 | 1.463657 | Upregulated |
| acute | olr1696 | 1.437133 | Upregulated |
| acute | mir382 | 1.42165 | Upregulated |
| acute | mir369 | 1.41 | Upregulated |
| acute | rgd1565058 | 1.395887 | Upregulated |
| acute | loc500990 | 1.384788 | Upregulated |
| acute | mir667 | 1.33 | Upregulated |
| acute | rgd1559532 | 1.318522 | Upregulated |
| acute | loc302576 | 1.312214 | Upregulated |
| acute | abo3 | 1.278143 | Upregulated |
| acute | loc689725 | 1.26751 | Upregulated |
| acute | rt1-m2 | 1.264793 | Upregulated |
| acute | glyatl1 | 1.260516 | Upregulated |
| acute | loc498933 | 1.250304 | Upregulated |
| acute | rt1-a3 | 1.247698 | Upregulated |
| acute | mir325 | 1.24 | Upregulated |
| acute | spink1l | 1.239981 | Upregulated |
| acute | rgd1564319 | 1.230984 | Upregulated |
| acute | mir29c | 1.23 | Upregulated |
| acute | clec4m | 1.216597 | Upregulated |
| acute | smim38 | 1.211639 | Upregulated |
| acute | rgd1565410 | 1.200117 | Upregulated |
| acute | best4 | 1.179689 | Upregulated |
| subacute | bnip5 | 2.98 | Upregulated |
| subacute | tmem121b | 2.85 | Upregulated |
| subacute | tafa1 | 2.68 | Upregulated |
| subacute | or4d1 | 2.64 | Upregulated |
| subacute | catspere2 | 2.49 | Upregulated |
| subacute | ldc1 | 1.96 | Upregulated |
| subacute | d430036j16rik | 1.94 | Upregulated |
| subacute | shisal2b | 1.88 | Upregulated |
| subacute | tafa2 | 1.86 | Upregulated |
| subacute | or51e2 | 1.84 | Upregulated |
| subacute | sfta3-ps | 1.82 | Upregulated |
| subacute | mup17 | 1.78 | Upregulated |
| subacute | nexmif | 1.76 | Upregulated |
| subacute | rtl4 | 1.72 | Upregulated |
| subacute | loc689064///hbb-b1 | 1.7 | Upregulated |
| subacute | gm10754 | 1.681 | Upregulated |
| subacute | cracdl | 1.67 | Upregulated |
| subacute | ccn3 | 1.586731 | Upregulated |
| subacute | entrep2 | 1.58 | Upregulated |
| subacute | gm10406 | 1.58 | Upregulated |
| subacute | lbhd2 | 1.572565 | Upregulated |
| subacute | or51a8 | 1.57 | Upregulated |
| subacute | trav3-3 | 1.565 | Upregulated |
| subacute | b230303a05rik | 1.551584 | Upregulated |
| subacute | mhrt | 1.5405 | Upregulated |
| subacute | gpr52 | 1.536 | Upregulated |
| subacute | gm3264 | 1.52 | Upregulated |
| subacute | or8k38 | 1.51 | Upregulated |
| subacute | or51f23 | 1.5 | Upregulated |
| subacute | gm10790 | 1.4881 | Upregulated |
| long-term | rgd1559482 | 5.706605 | Upregulated |
| long-term | msr1 | 5.189444 | Upregulated |
| long-term | cd300c2 | 5.144992 | Upregulated |
| long-term | stra6l | 4.002048 | Upregulated |
| long-term | cd300le | 3.975743 | Upregulated |
| long-term | lilrb2 | 3.70652 | Upregulated |
| long-term | ckap2l | 3.532761 | Upregulated |
| long-term | pclaf | 3.522151 | Upregulated |
| long-term | h1f5 | 3.145878 | Upregulated |
| long-term | calhm6 | 3.144975 | Upregulated |
| long-term | aoah | 3.143599 | Upregulated |
| long-term | rexo5 | 3.052205 | Upregulated |
| long-term | rt1-db2 | 2.935881 | Upregulated |
| long-term | ticam2 | 2.703448 | Upregulated |
| long-term | cd84 | 2.52246 | Upregulated |
| long-term | pimreg | 2.486643 | Upregulated |
| long-term | loc102553672 | 2.4358 | Upregulated |
| long-term | nlrp3 | 2.435346 | Upregulated |
| long-term | lrmda | 2.39649 | Upregulated |
| long-term | ugt1a2 | 2.339259 | Upregulated |
| long-term | loc102553978 | 2.28722 | Upregulated |
| long-term | loc689064///hbb-b1 | 2.26 | Upregulated |
| long-term | tmem273 | 2.224012 | Upregulated |
| long-term | loc102557492 | 2.22318 | Upregulated |
| long-term | c9h6orf141 | 2.196448 | Upregulated |
| long-term | tent5c | 2.102321 | Upregulated |
| long-term | tasl | 2.10042 | Upregulated |
| long-term | mastl | 2.074469 | Upregulated |
| long-term | kng1 | 2.04123 | Upregulated |
| long-term | rt1-bb | 2.035951 | Upregulated |
| acute | hspa1b | -3.38884 | Downregulated |
| acute | hspa1a | -3.24 | Downregulated |
| acute | npas4 | -3.10446 | Downregulated |
| acute | ccl3 | -2.93092 | Downregulated |
| acute | lilrb4a///lilr4b | -2.68959 | Downregulated |
| acute | pald1///thbs1 | -2.38141 | Downregulated |
| acute | fos | -2.30201 | Downregulated |
| acute | cxcl1 | -2.094 | Downregulated |
| acute | ccl2 | -2.08238 | Downregulated |
| acute | ccn1 | -2.00473 | Downregulated |
| acute | ctla2b///ctla2a | -1.97493 | Downregulated |
| acute | a130040m12rik | -1.96781 | Downregulated |
| acute | fosb | -1.92343 | Downregulated |
| acute | snord14e | -1.775 | Downregulated |
| acute | atf3 | -1.74609 | Downregulated |
| acute | a130040m12rik | -1.72056 | Downregulated |
| acute | cd14 | -1.67338 | Downregulated |
| acute | 3930401b19rik | -1.66604 | Downregulated |
| acute | loc100044068///ifi202b | -1.63828 | Downregulated |
| acute | apold1 | -1.61835 | Downregulated |
| acute | b230214o09rik | -1.57603 | Downregulated |
| acute | a130040m12rik | -1.55496 | Downregulated |
| acute | ccl4 | -1.53103 | Downregulated |
| acute | cyr61 | -1.47603 | Downregulated |
| acute | gadd45g | -1.41378 | Downregulated |
| acute | zfp36 | -1.37922 | Downregulated |
| acute | egr2 | -1.29178 | Downregulated |
| acute | hspb1 | -1.29 | Downregulated |
| acute | hmox1 | -1.2877 | Downregulated |
| acute | gadd45b | -1.21532 | Downregulated |
| subacute | lilrb4b | -7.22 | Downregulated |
| subacute | cstdc4 | -7.11 | Downregulated |
| subacute | pira12 | -6.19 | Downregulated |
| subacute | ifi211 | -5.95 | Downregulated |
| subacute | cstdc5 | -4.94 | Downregulated |
| subacute | pira2 | -4.935 | Downregulated |
| subacute | ifi207 | -4.875 | Downregulated |
| subacute | lilrb4a | -4.87 | Downregulated |
| subacute | ifi209 | -4.49 | Downregulated |
| subacute | rgd1559482 | -4.4486 | Downregulated |
| subacute | spp1 | -4.34252 | Downregulated |
| subacute | pira7 | -4.33 | Downregulated |
| subacute | gm3776 | -4.05 | Downregulated |
| subacute | lilrb4a///lilr4b | -3.92259 | Downregulated |
| subacute | mirt2 | -3.4273 | Downregulated |
| subacute | lcn2 | -3.40897 | Downregulated |
| subacute | lgals3 | -3.37851 | Downregulated |
| subacute | timp1 | -3.26356 | Downregulated |
| subacute | hspa1a | -3.03967 | Downregulated |
| subacute | sirpb1b | -2.97855 | Downregulated |
| subacute | gm9733 | -2.9735 | Downregulated |
| subacute | hspb1 | -2.97156 | Downregulated |
| subacute | ccl2 | -2.94708 | Downregulated |
| subacute | hmox1 | -2.92296 | Downregulated |
| subacute | pira11 | -2.87419 | Downregulated |
| subacute | ptprc///ptprc | -2.82 | Downregulated |
| subacute | ccl12 | -2.76286 | Downregulated |
| subacute | ccn1 | -2.75 | Downregulated |
| subacute | gm5150 | -2.7252 | Downregulated |
| subacute | jaml | -2.72 | Downregulated |
| long-term | ly49i4///ly49s7///klra5///ly49i2 | -6.19906 | Downregulated |
| long-term | clec7a | -5.44062 | Downregulated |
| long-term | rgd1561778 | -5.2629 | Downregulated |
| long-term | loc100910679 | -5.0462 | Downregulated |
| long-term | loc100911545///a2m | -4.63554 | Downregulated |
| long-term | kng1l1///kng2///kng1 | -4.63364 | Downregulated |
| long-term | tlr8 | -4.6066 | Downregulated |
| long-term | itgax | -4.59114 | Downregulated |
| long-term | cyp2d5///cyp2d1 | -4.5711 | Downregulated |
| long-term | evi2b | -4.55916 | Downregulated |
| long-term | loc102553715///mfap4 | -4.40692 | Downregulated |
| long-term | loc691173///ckap2 | -4.3528 | Downregulated |
| long-term | cenpe | -4.33966 | Downregulated |
| long-term | mki67 | -4.27138 | Downregulated |
| long-term | loc686151 | -4.19458 | Downregulated |
| long-term | loc100359539///rrm2 | -4.17038 | Downregulated |
| long-term | wdfy4 | -4.12 | Downregulated |
| long-term | loc100911825 | -4.0286 | Downregulated |
| long-term | loc100910669 | -3.76666 | Downregulated |
| long-term | ccl6 | -3.72159 | Downregulated |
| long-term | igkc | -3.6946 | Downregulated |
| long-term | spp1 | -3.67 | Downregulated |
| long-term | depdc1 | -3.65378 | Downregulated |
| long-term | gpnmb | -3.62183 | Downregulated |
| long-term | itih2 | -3.60034 | Downregulated |
| long-term | card14 | -3.52862 | Downregulated |
| long-term | lum | -3.49 | Downregulated |
| long-term | irf8 | -3.48909 | Downregulated |
| long-term | loc100911217 | -3.48422 | Downregulated |
| long-term | frrs1 | -3.47068 | Downregulated |

**Suppl. Table 8**: Top30 enriched gene ontology terms for biological process in rats compared to mice acutely after stroke

| **ID** | **Description** | **setSize** | **enrichmentScore** | **NES** | **pvalue** | **p.adjust** |
| --- | --- | --- | --- | --- | --- | --- |
| GO:0030855 | epithelial cell differentiation | 591 | 0.429999082 | 1.34374 | 1E-04 | 1E-04 |
| GO:0048598 | embryonic morphogenesis | 555 | 0.433473488 | 1.35337 | 0.0001 | 0.0001 |
| GO:0048732 | gland development | 411 | 0.482489403 | 1.497628 | 0.0001 | 0.0001 |
| GO:0045165 | cell fate commitment | 242 | 0.523522129 | 1.602086 | 0.0001 | 0.0001 |
| GO:0019953 | sexual reproduction | 680 | 0.409816569 | 1.284076 | 0.0003 | 0.0003 |
| GO:0098660 | inorganic ion transmembrane transport | 690 | 0.409200905 | 1.282466 | 0.0003 | 0.0003 |
| GO:0050954 | sensory perception of mechanical stimulus | 169 | 0.501744814 | 1.510789 | 0.000305 | 0.000305 |
| GO:0003006 | developmental process involved in reproduction | 761 | 0.406563244 | 1.275806 | 0.0004 | 0.0004 |
| GO:0048736 | appendage development | 160 | 0.506426564 | 1.519896 | 0.000509 | 0.000509 |
| GO:0060173 | limb development | 160 | 0.506426564 | 1.519896 | 0.000509 | 0.000509 |
| GO:0008544 | epidermis development | 283 | 0.455083699 | 1.399984 | 0.000601 | 0.000601 |
| GO:0050953 | sensory perception of light stimulus | 139 | 0.507268979 | 1.508509 | 0.00082 | 0.00082 |
| GO:0048706 | embryonic skeletal system development | 112 | 0.524257342 | 1.538371 | 0.001035 | 0.001035 |
| GO:0006936 | muscle contraction | 276 | 0.452288136 | 1.389658 | 0.001103 | 0.001103 |
| GO:0098739 | import across plasma membrane | 182 | 0.480666531 | 1.454108 | 0.001214 | 0.001214 |
| GO:0007009 | plasma membrane organization | 138 | 0.496957773 | 1.47718 | 0.001436 | 0.001436 |
| GO:0060041 | retina development in camera-type eye | 145 | 0.487725357 | 1.455216 | 0.001534 | 0.001534 |
| GO:0061061 | muscle structure development | 587 | 0.39911016 | 1.247152 | 0.0027 | 0.0027 |
| GO:0060485 | mesenchyme development | 266 | 0.444042247 | 1.363023 | 0.00281 | 0.00281 |
| GO:0022600 | digestive system process | 107 | 0.505476143 | 1.478246 | 0.003213 | 0.003213 |
| GO:0042692 | muscle cell differentiation | 363 | 0.420429475 | 1.302239 | 0.004003 | 0.004003 |
| GO:0050678 | regulation of epithelial cell proliferation | 372 | 0.417276208 | 1.292752 | 0.004103 | 0.004103 |
| GO:0045766 | positive regulation of angiogenesis | 158 | 0.469051833 | 1.406402 | 0.004179 | 0.004179 |
| GO:1904018 | positive regulation of vasculature development | 158 | 0.469051833 | 1.406402 | 0.004179 | 0.004179 |
| GO:0010876 | lipid localization | 397 | 0.416005842 | 1.290353 | 0.004404 | 0.004404 |
| GO:0008202 | steroid metabolic process | 279 | 0.433385326 | 1.332341 | 0.004612 | 0.004612 |
| GO:0030324 | lung development | 196 | 0.451858211 | 1.371244 | 0.004849 | 0.004849 |
| GO:0007043 | cell-cell junction assembly | 126 | 0.482893535 | 1.428081 | 0.004938 | 0.004938 |
| GO:0007586 | digestion | 122 | 0.485820212 | 1.433444 | 0.00495 | 0.00495 |
| GO:0045321 | leukocyte activation | 773 | 0.383951627 | 1.204933 | 0.005 | 0.005 |

**Suppl. Table 9:** Top30 enriched gene ontology terms for biological process in rats compared to mice subacutely after stroke

| **ID** | **Description** | **setSize** | **enrichmentScore** | **NES** | **pvalue** | **p.adjust** |
| --- | --- | --- | --- | --- | --- | --- |
| GO:0002684 | positive regulation of immune system process | 795 | -0.360076611 | -1.4397412 | 0.00010268 | 0.00010268 |
| GO:0046649 | lymphocyte activation | 716 | -0.370986694 | -1.4774637 | 0.00010349 | 0.00010349 |
| GO:0050776 | regulation of immune response | 709 | -0.377369353 | -1.5022166 | 0.00010356 | 0.00010356 |
| GO:1903047 | mitotic cell cycle process | 706 | -0.47248094 | -1.8809582 | 0.00010357 | 0.00010357 |
| GO:0010564 | regulation of cell cycle process | 663 | -0.413769365 | -1.6421426 | 0.0001041 | 0.0001041 |
| GO:0051301 | cell division | 604 | -0.432095013 | -1.7087828 | 0.0001047 | 0.0001047 |
| GO:0002694 | regulation of leukocyte activation | 554 | -0.377859516 | -1.4882504 | 0.00010546 | 0.00010546 |
| GO:0006281 | DNA repair | 523 | -0.400887482 | -1.5751859 | 0.00010594 | 0.00010594 |
| GO:0044770 | cell cycle phase transition | 511 | -0.460460636 | -1.8064658 | 0.00010632 | 0.00010632 |
| GO:0042110 | T cell activation | 501 | -0.379200183 | -1.4860331 | 0.00010659 | 0.00010659 |
| GO:0048285 | organelle fission | 474 | -0.46031201 | -1.7976881 | 0.00010749 | 0.00010749 |
| GO:0051276 | chromosome organization | 476 | -0.484749824 | -1.8930767 | 0.00010754 | 0.00010754 |
| GO:0007346 | regulation of mitotic cell cycle | 476 | -0.442904511 | -1.7296597 | 0.00010754 | 0.00010754 |
| GO:0051249 | regulation of lymphocyte activation | 448 | -0.386996162 | -1.5073716 | 0.00010805 | 0.00010805 |
| GO:0000280 | nuclear division | 424 | -0.48180331 | -1.8721465 | 0.0001085 | 0.0001085 |
| GO:0044772 | mitotic cell cycle phase transition | 429 | -0.469876232 | -1.8263593 | 0.00010854 | 0.00010854 |
| GO:1901987 | regulation of cell cycle phase transition | 404 | -0.475943968 | -1.8436215 | 0.0001094 | 0.0001094 |
| GO:0010639 | negative regulation of organelle organization | 358 | -0.40293074 | -1.5487714 | 0.00011112 | 0.00011112 |
| GO:0045786 | negative regulation of cell cycle | 353 | -0.438105618 | -1.6826113 | 0.00011116 | 0.00011116 |
| GO:0007059 | chromosome segregation | 333 | -0.519751708 | -1.9881839 | 0.00011174 | 0.00011174 |
| GO:1901990 | regulation of mitotic cell cycle phase transition | 323 | -0.483842929 | -1.8471584 | 0.00011196 | 0.00011196 |
| GO:0140014 | mitotic nuclear division | 289 | -0.484617064 | -1.8347469 | 0.00011337 | 0.00011337 |
| GO:0006310 | DNA recombination | 286 | -0.424011833 | -1.6041138 | 0.0001137 | 0.0001137 |
| GO:0098813 | nuclear chromosome segregation | 273 | -0.528824731 | -1.9945684 | 0.00011401 | 0.00011401 |
| GO:0010948 | negative regulation of cell cycle process | 279 | -0.485187993 | -1.8308539 | 0.00011419 | 0.00011419 |
| GO:0006302 | double-strand break repair | 270 | -0.434014191 | -1.6349364 | 0.00011439 | 0.00011439 |
| GO:0006260 | DNA replication | 262 | -0.448000861 | -1.6823588 | 0.00011494 | 0.00011494 |
| GO:0090068 | positive regulation of cell cycle process | 242 | -0.453882621 | -1.6955851 | 0.00011606 | 0.00011606 |
| GO:0051321 | meiotic cell cycle | 241 | -0.46309468 | -1.7291819 | 0.00011623 | 0.00011623 |
| GO:1901988 | negative regulation of cell cycle phase transition | 238 | -0.508342548 | -1.8960014 | 0.00011641 | 0.00011641 |

**Suppl. Table 10**: Top30 enriched gene ontology terms for biological process in rats compared to mice long-term after stroke

| **ID** | **Description** | **setSize** | **enrichmentScore** | **NES** | **pvalue** | **p.adjust** |
| --- | --- | --- | --- | --- | --- | --- |
| GO:0099536 | synaptic signaling | 697 | 0.429860312 | 1.99335644 | 0.00015181 | 0.00015181 |
| GO:0007610 | behavior | 650 | 0.358753328 | 1.6550548 | 0.00015323 | 0.00015323 |
| GO:0043269 | regulation of ion transport | 640 | 0.333568297 | 1.53733107 | 0.00015326 | 0.00015326 |
| GO:0007186 | G protein-coupled receptor signaling pathway | 529 | 0.378830028 | 1.71918034 | 0.00015593 | 0.00015593 |
| GO:0099177 | regulation of trans-synaptic signaling | 485 | 0.444569588 | 2.00158813 | 0.00015669 | 0.00015669 |
| GO:0050804 | modulation of chemical synaptic transmission | 484 | 0.44465655 | 2.00152376 | 0.00015669 | 0.00015669 |
| GO:0050808 | synapse organization | 469 | 0.352465811 | 1.58195599 | 0.00015798 | 0.00015798 |
| GO:0006887 | exocytosis | 346 | 0.394600063 | 1.72204948 | 0.00016197 | 0.00016197 |
| GO:0050890 | cognition | 325 | 0.388615127 | 1.68642291 | 0.00016236 | 0.00016236 |
| GO:0099003 | vesicle-mediated transport in synapse | 246 | 0.384593668 | 1.62249697 | 0.00016628 | 0.00016628 |
| GO:0016079 | synaptic vesicle exocytosis | 129 | 0.458239582 | 1.77775857 | 0.00017232 | 0.00017232 |
| GO:0002761 | regulation of myeloid leukocyte differentiation | 106 | -0.44679175 | -1.7434698 | 0.00023408 | 0.00023408 |
| GO:0034446 | substrate adhesion-dependent cell spreading | 104 | -0.470403856 | -1.829429 | 0.00023419 | 0.00023419 |
| GO:0030183 | B cell differentiation | 113 | -0.478191449 | -1.8869574 | 0.0002354 | 0.0002354 |
| GO:0043534 | blood vessel endothelial cell migration | 112 | -0.469042447 | -1.8463886 | 0.00023568 | 0.00023568 |
| GO:0072089 | stem cell proliferation | 114 | -0.44139276 | -1.7428473 | 0.00023697 | 0.00023697 |
| GO:0051304 | chromosome separation | 114 | -0.437878194 | -1.72897 | 0.00023697 | 0.00023697 |
| GO:0050817 | coagulation | 132 | -0.468612618 | -1.8926014 | 0.00023958 | 0.00023958 |
| GO:0007599 | hemostasis | 132 | -0.453480152 | -1.8314855 | 0.00023958 | 0.00023958 |
| GO:0002262 | myeloid cell homeostasis | 161 | -0.45595278 | -1.8987019 | 0.00024266 | 0.00024266 |
| GO:0042113 | B cell activation | 211 | -0.412696453 | -1.7781665 | 0.00024426 | 0.00024426 |
| GO:0022409 | positive regulation of cell-cell adhesion | 230 | -0.383594472 | -1.667135 | 0.00024814 | 0.00024814 |
| GO:0060326 | cell chemotaxis | 226 | -0.368374992 | -1.5980327 | 0.00024839 | 0.00024839 |
| GO:0003007 | heart morphogenesis | 234 | -0.357673084 | -1.5579432 | 0.00024876 | 0.00024876 |
| GO:0051251 | positive regulation of lymphocyte activation | 236 | -0.363351968 | -1.5841191 | 0.00024913 | 0.00024913 |
| GO:1901342 | regulation of vasculature development | 253 | -0.368141339 | -1.6199137 | 0.000251 | 0.000251 |
| GO:0070661 | leukocyte proliferation | 260 | -0.357555605 | -1.57769 | 0.00025132 | 0.00025132 |
| GO:0002764 | immune response-regulating signaling pathway | 276 | -0.384999442 | -1.7090475 | 0.00025151 | 0.00025151 |
| GO:0090132 | epithelium migration | 285 | -0.382095543 | -1.7017339 | 0.00025374 | 0.00025374 |
| GO:0042060 | wound healing | 301 | -0.360506963 | -1.6151513 | 0.0002551 | 0.0002551 |

**Suppl. Table 11**: Top30 enriched gene ontology terms for biological process in pMCAo compared to tMCAo acutely after stroke

| **ID** | **Description** | **setSize** | **enrichmentScore** | **NES** | **pvalue** | **p.adjust** |
| --- | --- | --- | --- | --- | --- | --- |
| GO:0050906 | detection of stimulus involved in sensory perception | 176 | -0.461438564 | -1.84255 | 0.000158 | 0.000158 |
| GO:0009593 | detection of chemical stimulus | 125 | -0.492701337 | -1.88177 | 0.000164 | 0.000164 |
| GO:0050829 | defense response to Gram-negative bacterium | 113 | -0.512371378 | -1.93068 | 0.000166 | 0.000166 |
| GO:0007229 | integrin-mediated signaling pathway | 103 | 0.460995975 | 1.805303 | 0.000248 | 0.000248 |
| GO:0060993 | kidney morphogenesis | 104 | 0.463711933 | 1.820313 | 0.000248 | 0.000248 |
| GO:0070167 | regulation of biomineral tissue development | 107 | 0.474148612 | 1.867557 | 0.000249 | 0.000249 |
| GO:0061326 | renal tubule development | 108 | 0.461186368 | 1.820016 | 0.000249 | 0.000249 |
| GO:0016525 | negative regulation of angiogenesis | 108 | 0.458536507 | 1.809558 | 0.000249 | 0.000249 |
| GO:0010657 | muscle cell apoptotic process | 108 | 0.445240227 | 1.757086 | 0.000249 | 0.000249 |
| GO:0034446 | substrate adhesion-dependent cell spreading | 109 | 0.498623966 | 1.971194 | 0.00025 | 0.00025 |
| GO:0110149 | regulation of biomineralization | 109 | 0.465084712 | 1.838605 | 0.00025 | 0.00025 |
| GO:0048661 | positive regulation of smooth muscle cell proliferation | 109 | 0.438750695 | 1.734499 | 0.00025 | 0.00025 |
| GO:0071346 | cellular response to interferon-gamma | 110 | 0.495438194 | 1.962478 | 0.00025 | 0.00025 |
| GO:2000181 | negative regulation of blood vessel morphogenesis | 110 | 0.455140703 | 1.802855 | 0.00025 | 0.00025 |
| GO:0110020 | regulation of actomyosin structure organization | 110 | 0.432895877 | 1.714742 | 0.00025 | 0.00025 |
| GO:1901343 | negative regulation of vasculature development | 111 | 0.453550338 | 1.797774 | 0.000251 | 0.000251 |
| GO:0008630 | intrinsic apoptotic signaling pathway in response to DNA damage | 113 | 0.448254505 | 1.780878 | 0.000253 | 0.000253 |
| GO:0021987 | cerebral cortex development | 113 | 0.447427145 | 1.777591 | 0.000253 | 0.000253 |
| GO:0010769 | regulation of cell morphogenesis involved in differentiation | 121 | 0.457358154 | 1.839851 | 0.000253 | 0.000253 |
| GO:0003279 | cardiac septum development | 122 | 0.473824296 | 1.907483 | 0.000254 | 0.000254 |
| GO:0048709 | oligodendrocyte differentiation | 114 | 0.508908526 | 2.023963 | 0.000254 | 0.000254 |
| GO:0060840 | artery development | 114 | 0.48198422 | 1.916883 | 0.000254 | 0.000254 |
| GO:0032231 | regulation of actin filament bundle assembly | 114 | 0.44526826 | 1.770861 | 0.000254 | 0.000254 |
| GO:0019079 | viral genome replication | 123 | 0.435637413 | 1.756397 | 0.000254 | 0.000254 |
| GO:1901989 | positive regulation of cell cycle phase transition | 115 | 0.495165502 | 1.971214 | 0.000254 | 0.000254 |
| GO:0045446 | endothelial cell differentiation | 119 | 0.487067315 | 1.95336 | 0.000254 | 0.000254 |
| GO:0021782 | glial cell development | 120 | 0.478081034 | 1.920468 | 0.000255 | 0.000255 |
| GO:0016241 | regulation of macroautophagy | 117 | 0.47303153 | 1.890746 | 0.000255 | 0.000255 |
| GO:0072009 | nephron epithelium development | 117 | 0.442981118 | 1.770632 | 0.000255 | 0.000255 |
| GO:0048565 | digestive tract development | 124 | 0.43046716 | 1.736576 | 0.000255 | 0.000255 |

**Suppl. Table 12**: Top30 enriched gene ontology terms for biological process in pMCAo compared to tMCAo long-term after stroke

| **ID** | **Description** | **setSize** | **enrichmentScore** | **NES** | **pvalue** | **p.adjust** |
| --- | --- | --- | --- | --- | --- | --- |
| GO:0098542 | defense response to other organism | 474 | 0.696883519 | 2.22599414 | 9.999E-05 | 9.999E-05 |
| GO:0045087 | innate immune response | 349 | 0.693042705 | 2.20348366 | 9.999E-05 | 9.999E-05 |
| GO:0050776 | regulation of immune response | 379 | 0.687647182 | 2.19006944 | 9.999E-05 | 9.999E-05 |
| GO:0006954 | inflammatory response | 393 | 0.684514254 | 2.18105576 | 9.999E-05 | 9.999E-05 |
| GO:0006955 | immune response | 714 | 0.678030912 | 2.17735017 | 9.999E-05 | 9.999E-05 |
| GO:0051707 | response to other organism | 618 | 0.675969629 | 2.16773917 | 9.999E-05 | 9.999E-05 |
| GO:0043207 | response to external biotic stimulus | 619 | 0.674932699 | 2.1644106 | 9.999E-05 | 9.999E-05 |
| GO:0002684 | positive regulation of immune system process | 450 | 0.677608944 | 2.16187552 | 9.999E-05 | 9.999E-05 |
| GO:0009607 | response to biotic stimulus | 641 | 0.669754695 | 2.14902139 | 9.999E-05 | 9.999E-05 |
| GO:0044419 | biological process involved in interspecies interaction between organisms | 688 | 0.665447954 | 2.13696658 | 9.999E-05 | 9.999E-05 |
| GO:0006952 | defense response | 767 | 0.662238587 | 2.1288953 | 9.999E-05 | 9.999E-05 |
| GO:0002682 | regulation of immune system process | 720 | 0.651710263 | 2.09284381 | 9.999E-05 | 9.999E-05 |
| GO:0046649 | lymphocyte activation | 391 | 0.654866341 | 2.08688276 | 9.999E-05 | 9.999E-05 |
| GO:0045321 | leukocyte activation | 486 | 0.646912568 | 2.06717925 | 9.999E-05 | 9.999E-05 |
| GO:0050865 | regulation of cell activation | 340 | 0.646942707 | 2.05574691 | 9.999E-05 | 9.999E-05 |
| GO:0001775 | cell activation | 553 | 0.642004721 | 2.0548027 | 9.999E-05 | 9.999E-05 |
| GO:0001816 | cytokine production | 416 | 0.642656009 | 2.04940683 | 9.999E-05 | 9.999E-05 |
| GO:0001817 | regulation of cytokine production | 380 | 0.641684046 | 2.04388451 | 9.999E-05 | 9.999E-05 |
| GO:0032101 | regulation of response to external stimulus | 512 | 0.613064653 | 1.96041729 | 9.999E-05 | 9.999E-05 |
| GO:0030335 | positive regulation of cell migration | 350 | 0.603206478 | 1.91811218 | 9.999E-05 | 9.999E-05 |
| GO:0034097 | response to cytokine | 457 | 0.596586144 | 1.90421268 | 9.999E-05 | 9.999E-05 |
| GO:2000147 | positive regulation of cell motility | 361 | 0.595395862 | 1.89356146 | 9.999E-05 | 9.999E-05 |
| GO:0040017 | positive regulation of locomotion | 370 | 0.59010494 | 1.87809613 | 9.999E-05 | 9.999E-05 |
| GO:0071345 | cellular response to cytokine stimulus | 403 | 0.587803933 | 1.87341178 | 9.999E-05 | 9.999E-05 |
| GO:0030155 | regulation of cell adhesion | 442 | 0.586760681 | 1.87204523 | 9.999E-05 | 9.999E-05 |
| GO:0002520 | immune system development | 542 | 0.579305333 | 1.85360992 | 9.999E-05 | 9.999E-05 |
| GO:0048534 | hematopoietic or lymphoid organ development | 518 | 0.575060911 | 1.8389758 | 9.999E-05 | 9.999E-05 |
| GO:0030097 | hemopoiesis | 498 | 0.575047195 | 1.83820796 | 9.999E-05 | 9.999E-05 |
| GO:0030334 | regulation of cell migration | 583 | 0.556474862 | 1.78305074 | 9.999E-05 | 9.999E-05 |
| GO:2000145 | regulation of cell motility | 609 | 0.548304192 | 1.75791002 | 9.999E-05 | 9.999E-05 |

**Suppl. Table 13**: Top30 enriched gene ontology terms for biological process in PT compared to tMCAo long-term after stroke

| **ID** | **Description** | **setSize** | **enrichmentScore** | **NES** | **pvalue** | **p.adjust** |
| --- | --- | --- | --- | --- | --- | --- |
| GO:0008284 | positive regulation of cell population proliferation | 795 | 0.349142081 | 1.59128507 | 0.00012271 | 0.00012271 |
| GO:0000278 | mitotic cell cycle | 799 | 0.373994051 | 1.70478078 | 0.00012273 | 0.00012273 |
| GO:0048534 | hematopoietic or lymphoid organ development | 792 | 0.425197764 | 1.9366533 | 0.00012308 | 0.00012308 |
| GO:0043066 | negative regulation of apoptotic process | 788 | 0.313436538 | 1.42698109 | 0.00012337 | 0.00012337 |
| GO:0051336 | regulation of hydrolase activity | 765 | 0.343797118 | 1.56336549 | 0.00012346 | 0.00012346 |
| GO:0030097 | hemopoiesis | 759 | 0.424221791 | 1.92716081 | 0.00012379 | 0.00012379 |
| GO:0032101 | regulation of response to external stimulus | 720 | 0.395471769 | 1.79025172 | 0.00012513 | 0.00012513 |
| GO:0045321 | leukocyte activation | 714 | 0.425825598 | 1.92670129 | 0.00012525 | 0.00012525 |
| GO:0030029 | actin filament-based process | 711 | 0.338863762 | 1.53260209 | 0.0001255 | 0.0001255 |
| GO:0009792 | embryo development ending in birth or egg hatching | 705 | 0.322717082 | 1.45902512 | 0.00012552 | 0.00012552 |
| GO:0043009 | chordate embryonic development | 692 | 0.327699162 | 1.47969331 | 0.0001259 | 0.0001259 |
| GO:0098609 | cell-cell adhesion | 679 | 0.384706693 | 1.73573447 | 0.00012623 | 0.00012623 |
| GO:0001944 | vasculature development | 674 | 0.435117363 | 1.96233565 | 0.00012633 | 0.00012633 |
| GO:1903047 | mitotic cell cycle process | 675 | 0.402327961 | 1.8146799 | 0.00012633 | 0.00012633 |
| GO:0030155 | regulation of cell adhesion | 653 | 0.405560865 | 1.82597772 | 0.00012653 | 0.00012653 |
| GO:0034097 | response to cytokine | 656 | 0.421760861 | 1.89955621 | 0.00012658 | 0.00012658 |
| GO:0098542 | defense response to other organism | 658 | 0.438600053 | 1.97512402 | 0.00012677 | 0.00012677 |
| GO:0002684 | positive regulation of immune system process | 630 | 0.461125589 | 2.07116311 | 0.00012708 | 0.00012708 |
| GO:0001568 | blood vessel development | 641 | 0.439556721 | 1.97503984 | 0.00012723 | 0.00012723 |
| GO:0030036 | actin cytoskeleton organization | 634 | 0.35649769 | 1.60098394 | 0.00012726 | 0.00012726 |
| GO:0010564 | regulation of cell cycle process | 620 | 0.376634312 | 1.68884583 | 0.0001277 | 0.0001277 |
| GO:0008285 | negative regulation of cell population proliferation | 602 | 0.334582564 | 1.49736034 | 0.00012845 | 0.00012845 |
| GO:0046649 | lymphocyte activation | 591 | 0.437313005 | 1.95429605 | 0.00012882 | 0.00012882 |
| GO:0010942 | positive regulation of cell death | 591 | 0.351136624 | 1.56918479 | 0.00012882 | 0.00012882 |
| GO:0001816 | cytokine production | 595 | 0.450401907 | 2.01305322 | 0.00012907 | 0.00012907 |
| GO:0071345 | cellular response to cytokine stimulus | 589 | 0.430686796 | 1.92392512 | 0.0001292 | 0.0001292 |
| GO:0048729 | tissue morphogenesis | 573 | 0.353331304 | 1.57552899 | 0.00012968 | 0.00012968 |
| GO:0033993 | response to lipid | 540 | 0.346713417 | 1.54153539 | 0.00012999 | 0.00012999 |
| GO:0048514 | blood vessel morphogenesis | 561 | 0.458761833 | 2.04262082 | 0.00013004 | 0.00013004 |
| GO:0001817 | regulation of cytokine production | 543 | 0.453146639 | 2.01461302 | 0.00013012 | 0.00013012 |

**Suppl. Table 14**: Top30 enriched gene ontology terms for biological process in PT compared to pMCAo long-term after stroke

| **ID** | **Description** | **setSize** | **enrichmentScore** | **NES** | **pvalue** | **p.adjust** |
| --- | --- | --- | --- | --- | --- | --- |
| GO:0098542 | defense response to other organism | 423 | -0.7042271 | -2.1723258 | 9.999E-05 | 9.999E-05 |
| GO:0050776 | regulation of immune response | 361 | -0.6962554 | -2.1429945 | 9.999E-05 | 9.999E-05 |
| GO:0006955 | immune response | 648 | -0.6852093 | -2.1217268 | 9.999E-05 | 9.999E-05 |
| GO:0051707 | response to other organism | 560 | -0.6831594 | -2.1117218 | 9.999E-05 | 9.999E-05 |
| GO:0043207 | response to external biotic stimulus | 561 | -0.6820706 | -2.1084829 | 9.999E-05 | 9.999E-05 |
| GO:0009607 | response to biotic stimulus | 581 | -0.6764956 | -2.0921311 | 9.999E-05 | 9.999E-05 |
| GO:0044419 | biological process involved in interaction between organisms | 627 | -0.6735405 | -2.0844504 | 9.999E-05 | 9.999E-05 |
| GO:0006952 | defense response | 694 | -0.6682557 | -2.0697583 | 9.999E-05 | 9.999E-05 |
| GO:0002684 | positive regulation of immune system process | 421 | -0.6680067 | -2.0604866 | 9.999E-05 | 9.999E-05 |
| GO:0045321 | leukocyte activation | 460 | -0.6517877 | -2.0113865 | 9.999E-05 | 9.999E-05 |
| GO:0002682 | regulation of immune system process | 675 | -0.6468376 | -2.0029946 | 9.999E-05 | 9.999E-05 |
| GO:0001817 | regulation of cytokine production | 360 | -0.6497896 | -1.9998643 | 9.999E-05 | 9.999E-05 |
| GO:0046649 | lymphocyte activation | 372 | -0.6473968 | -1.9935965 | 9.999E-05 | 9.999E-05 |
| GO:0001816 | cytokine production | 392 | -0.6464315 | -1.992206 | 9.999E-05 | 9.999E-05 |
| GO:0001775 | cell activation | 523 | -0.6427989 | -1.9855595 | 9.999E-05 | 9.999E-05 |
| GO:0032101 | regulation of response to external stimulus | 474 | -0.6156012 | -1.9000375 | 9.999E-05 | 9.999E-05 |
| GO:0034097 | response to cytokine | 440 | -0.5957394 | -1.8381373 | 9.999E-05 | 9.999E-05 |
| GO:0071345 | cellular response to cytokine stimulus | 390 | -0.5892119 | -1.8154751 | 9.999E-05 | 9.999E-05 |
| GO:0030155 | regulation of cell adhesion | 421 | -0.5878985 | -1.8133906 | 9.999E-05 | 9.999E-05 |
| GO:0002520 | immune system development | 519 | -0.5664523 | -1.7497155 | 9.999E-05 | 9.999E-05 |
| GO:0030097 | hemopoiesis | 476 | -0.565256 | -1.7447638 | 9.999E-05 | 9.999E-05 |
| GO:0048534 | hematopoietic or lymphoid organ development | 495 | -0.5628788 | -1.7380183 | 9.999E-05 | 9.999E-05 |
| GO:0030334 | regulation of cell migration | 557 | -0.5519719 | -1.7062962 | 9.999E-05 | 9.999E-05 |
| GO:0048514 | blood vessel morphogenesis | 382 | -0.5534341 | -1.7047362 | 9.999E-05 | 9.999E-05 |
| GO:2000145 | regulation of cell motility | 579 | -0.5480803 | -1.6950066 | 9.999E-05 | 9.999E-05 |
| GO:0010628 | positive regulation of gene expression | 569 | -0.5409359 | -1.6721705 | 9.999E-05 | 9.999E-05 |
| GO:0080134 | regulation of response to stress | 729 | -0.5361915 | -1.6621738 | 9.999E-05 | 9.999E-05 |
| GO:0040012 | regulation of locomotion | 609 | -0.5360312 | -1.6586828 | 9.999E-05 | 9.999E-05 |
| GO:0033993 | response to lipid | 359 | -0.5365658 | -1.6512435 | 9.999E-05 | 9.999E-05 |
| GO:0001568 | blood vessel development | 436 | -0.5312964 | -1.6393371 | 9.999E-05 | 9.999E-05 |

**Suppl. Table 15:** Top30 enriched gene ontology terms for biological process in long-term compared to acute stroke.

| **ID** | **Description** | **setSize** | **enrichmentScore** | **NES** | **pvalue** | **p.adjust** |
| --- | --- | --- | --- | --- | --- | --- |
| GO:0009607 | response to biotic stimulus | 756 | -0.376647591 | -1.56524 | 0.000104 | 0.000104 |
| GO:0043207 | response to external biotic stimulus | 729 | -0.377960361 | -1.56843 | 0.000104 | 0.000104 |
| GO:0051707 | response to other organism | 729 | -0.377960361 | -1.56843 | 0.000104 | 0.000104 |
| GO:0001775 | cell activation | 620 | -0.387725161 | -1.59772 | 0.000106 | 0.000106 |
| GO:0098542 | defense response to other organism | 548 | -0.424258713 | -1.73921 | 0.000107 | 0.000107 |
| GO:0045321 | leukocyte activation | 544 | -0.387178801 | -1.58653 | 0.000107 | 0.000107 |
| GO:0002684 | positive regulation of immune system process | 500 | -0.389218104 | -1.58771 | 0.000108 | 0.000108 |
| GO:0046649 | lymphocyte activation | 453 | -0.392510951 | -1.59108 | 0.000109 | 0.000109 |
| GO:0050776 | regulation of immune response | 451 | -0.420215378 | -1.70275 | 0.000109 | 0.000109 |
| GO:0045087 | innate immune response | 394 | -0.451302939 | -1.81326 | 0.000111 | 0.000111 |
| GO:0002252 | immune effector process | 339 | -0.426574693 | -1.69539 | 0.000114 | 0.000114 |
| GO:0002694 | regulation of leukocyte activation | 332 | -0.398245405 | -1.57959 | 0.000114 | 0.000114 |
| GO:0050778 | positive regulation of immune response | 289 | -0.425740027 | -1.67078 | 0.000116 | 0.000116 |
| GO:0002250 | adaptive immune response | 230 | -0.459440049 | -1.76572 | 0.00012 | 0.00012 |
| GO:0050867 | positive regulation of cell activation | 223 | -0.424828246 | -1.62736 | 0.00012 | 0.00012 |
| GO:0002764 | immune response-regulating signaling pathway | 217 | -0.459280246 | -1.75572 | 0.000121 | 0.000121 |
| GO:0002443 | leukocyte mediated immunity | 214 | -0.4459233 | -1.70093 | 0.000121 | 0.000121 |
| GO:0070661 | leukocyte proliferation | 207 | -0.448560532 | -1.70639 | 0.000122 | 0.000122 |
| GO:0002696 | positive regulation of leukocyte activation | 207 | -0.43570102 | -1.65747 | 0.000122 | 0.000122 |
| GO:0009615 | response to virus | 184 | -0.452479748 | -1.7029 | 0.000123 | 0.000123 |
| GO:0002253 | activation of immune response | 178 | -0.454970573 | -1.70666 | 0.000124 | 0.000124 |
| GO:0046651 | lymphocyte proliferation | 181 | -0.447219091 | -1.67972 | 0.000124 | 0.000124 |
| GO:0042113 | B cell activation | 172 | -0.475909416 | -1.77915 | 0.000124 | 0.000124 |
| GO:0002449 | lymphocyte mediated immunity | 160 | -0.479157657 | -1.7793 | 0.000126 | 0.000126 |
| GO:0002460 | adaptive immune response based on somatic recombination…. | 161 | -0.497772441 | -1.84913 | 0.000126 | 0.000126 |
| GO:0002263 | cell activation involved in immune response | 156 | -0.465130034 | -1.72103 | 0.000126 | 0.000126 |
| GO:0001818 | negative regulation of cytokine production | 153 | -0.489233392 | -1.80412 | 0.000126 | 0.000126 |
| GO:0070663 | regulation of leukocyte proliferation | 153 | -0.483725348 | -1.78381 | 0.000126 | 0.000126 |
| GO:0002366 | leukocyte activation involved in immune response | 153 | -0.468340404 | -1.72708 | 0.000126 | 0.000126 |
| GO:0051607 | defense response to virus | 153 | -0.458299087 | -1.69005 | 0.000126 | 0.000126 |

**Suppl. Table 16**: Top30 enriched gene ontology terms for biological process in subacute to acute stroke.

| **ID** | **Description** | **setSize** | **enrichmentScore** | **NES** | **pvalue** | **p.adjust** |
| --- | --- | --- | --- | --- | --- | --- |
| GO:0007268 | chemical synaptic transmission | 787 | 0.393816226 | 1.61891102 | 0.00012123 | 0.00012123 |
| GO:0098916 | anterograde trans-synaptic signaling | 787 | 0.393816226 | 1.61891102 | 0.00012123 | 0.00012123 |
| GO:0099537 | trans-synaptic signaling | 795 | 0.394838615 | 1.62324161 | 0.00012134 | 0.00012134 |
| GO:0007610 | behavior | 766 | 0.381981351 | 1.56802026 | 0.00012154 | 0.00012154 |
| GO:0032990 | cell part morphogenesis | 747 | 0.359022921 | 1.47135665 | 0.00012209 | 0.00012209 |
| GO:0048858 | cell projection morphogenesis | 724 | 0.361270546 | 1.47782626 | 0.00012299 | 0.00012299 |
| GO:0120039 | plasma membrane bounded cell projection morphogenesis | 719 | 0.370053809 | 1.5131953 | 0.00012311 | 0.00012311 |
| GO:0048812 | neuron projection morphogenesis | 704 | 0.381615151 | 1.55729646 | 0.00012379 | 0.00012379 |
| GO:0048667 | cell morphogenesis involved in neuron differentiation | 654 | 0.381207356 | 1.55009213 | 0.00012492 | 0.00012492 |
| GO:0050804 | modulation of chemical synaptic transmission | 571 | 0.379099837 | 1.52852958 | 0.00012758 | 0.00012758 |
| GO:0099177 | regulation of trans-synaptic signaling | 572 | 0.378781459 | 1.52724537 | 0.00012776 | 0.00012776 |
| GO:0050808 | synapse organization | 493 | 0.430084867 | 1.71723163 | 0.00013004 | 0.00013004 |
| GO:0007611 | learning or memory | 319 | 0.426204918 | 1.6456454 | 0.00013805 | 0.00013805 |
| GO:0016358 | dendrite development | 309 | 0.409687304 | 1.57700581 | 0.00013856 | 0.00013856 |
| GO:0050803 | regulation of synapse structure or activity | 271 | 0.448317604 | 1.70782096 | 0.00014035 | 0.00014035 |
| GO:0050807 | regulation of synapse organization | 264 | 0.447376667 | 1.69862848 | 0.00014073 | 0.00014073 |
| GO:0099003 | vesicle-mediated transport in synapse | 259 | 0.46762199 | 1.77172142 | 0.00014104 | 0.00014104 |
| GO:0007626 | locomotory behavior | 262 | 0.450338933 | 1.70765428 | 0.00014116 | 0.00014116 |
| GO:0006836 | neurotransmitter transport | 242 | 0.439135106 | 1.65224518 | 0.00014255 | 0.00014255 |
| GO:0099504 | synaptic vesicle cycle | 229 | 0.455051808 | 1.70169647 | 0.00014418 | 0.00014418 |
| GO:0007416 | synapse assembly | 192 | 0.531770739 | 1.9528564 | 0.00014667 | 0.00014667 |
| GO:0007612 | learning | 186 | 0.495142018 | 1.80939565 | 0.0001481 | 0.0001481 |
| GO:0007269 | neurotransmitter secretion | 186 | 0.477126654 | 1.74356217 | 0.0001481 | 0.0001481 |
| GO:0099643 | signal release from synapse | 186 | 0.477126654 | 1.74356217 | 0.0001481 | 0.0001481 |
| GO:0048813 | dendrite morphogenesis | 185 | 0.453823847 | 1.65776341 | 0.0001481 | 0.0001481 |
| GO:0060078 | regulation of postsynaptic membrane potential | 117 | 0.556036126 | 1.92053776 | 0.00015375 | 0.00015375 |
| GO:0034330 | cell junction organization | 761 | 0.340323754 | 1.39656629 | 0.00024328 | 0.00024328 |
| GO:0001505 | regulation of neurotransmitter levels | 255 | 0.417925411 | 1.58195161 | 0.00028213 | 0.00028213 |
| GO:0032602 | chemokine production | 116 | -0.529397214 | -1.929309 | 0.00028289 | 0.00028289 |
| GO:0032755 | positive regulation of interleukin-6 production | 106 | -0.573818365 | -2.0560067 | 0.00028297 | 0.00028297 |

**Suppl. Table 17:** Top30 enriched gene ontology terms for biological process in long-term compared to subacute stroke.

| **ID** | **Description** | **setSize** | **enrichmentScore** | **NES** | **pvalue** | **p.adjust** |
| --- | --- | --- | --- | --- | --- | --- |
| GO:0006812 | cation transport | 742 | -0.355223167 | -1.3941575 | 0.00010854 | 0.00010854 |
| GO:0034220 | ion transmembrane transport | 627 | -0.394974776 | -1.5369613 | 0.00011073 | 0.00011073 |
| GO:0099536 | synaptic signaling | 540 | -0.473674841 | -1.8284524 | 0.00011222 | 0.00011222 |
| GO:0007610 | behavior | 513 | -0.401413109 | -1.5439399 | 0.00011296 | 0.00011296 |
| GO:0099537 | trans-synaptic signaling | 517 | -0.485003868 | -1.8658041 | 0.00011302 | 0.00011302 |
| GO:0007268 | chemical synaptic transmission | 510 | -0.486609707 | -1.8709924 | 0.00011305 | 0.00011305 |
| GO:0098916 | anterograde trans-synaptic signaling | 510 | -0.486609707 | -1.8709924 | 0.00011305 | 0.00011305 |
| GO:0098660 | inorganic ion transmembrane transport | 495 | -0.401517101 | -1.5411544 | 0.00011362 | 0.00011362 |
| GO:0043269 | regulation of ion transport | 487 | -0.399205434 | -1.530384 | 0.0001141 | 0.0001141 |
| GO:0099177 | regulation of trans-synaptic signaling | 372 | -0.491431402 | -1.8507284 | 0.00011816 | 0.00011816 |
| GO:0050804 | modulation of chemical synaptic transmission | 371 | -0.491694166 | -1.8509866 | 0.00011837 | 0.00011837 |
| GO:0034765 | regulation of ion transmembrane transport | 339 | -0.421329435 | -1.5765696 | 0.00011913 | 0.00011913 |
| GO:0042391 | regulation of membrane potential | 311 | -0.492472891 | -1.8288659 | 0.0001206 | 0.0001206 |
| GO:0050890 | cognition | 253 | -0.455451644 | -1.6621483 | 0.00012381 | 0.00012381 |
| GO:0007611 | learning or memory | 229 | -0.475828003 | -1.7198643 | 0.00012539 | 0.00012539 |
| GO:0048167 | regulation of synaptic plasticity | 166 | -0.525568258 | -1.8439742 | 0.00012999 | 0.00012999 |
| GO:0001508 | action potential | 107 | -0.593038137 | -1.9658162 | 0.0001385 | 0.0001385 |
| GO:0030001 | metal ion transport | 579 | -0.370988547 | -1.436044 | 0.00022411 | 0.00022411 |
| GO:0032989 | cellular component morphogenesis | 575 | -0.366882709 | -1.4198418 | 0.00022419 | 0.00022419 |
| GO:0098655 | cation transmembrane transport | 520 | -0.390691962 | -1.5039536 | 0.00022563 | 0.00022563 |
| GO:0032990 | cell part morphogenesis | 514 | -0.389037552 | -1.4963714 | 0.00022601 | 0.00022601 |
| GO:0120039 | plasma membrane bounded cell projection morphogenesis | 501 | -0.389421641 | -1.4959023 | 0.00022663 | 0.00022663 |
| GO:0048858 | cell projection morphogenesis | 503 | -0.390000315 | -1.4982277 | 0.00022665 | 0.00022665 |
| GO:0048812 | neuron projection morphogenesis | 491 | -0.395590416 | -1.5171437 | 0.00022774 | 0.00022774 |
| GO:0098662 | inorganic cation transmembrane transport | 455 | -0.394313979 | -1.5049811 | 0.00023052 | 0.00023052 |
| GO:0048667 | cell morphogenesis involved in neuron differentiation | 451 | -0.392657306 | -1.4977983 | 0.00023052 | 0.00023052 |
| GO:0007600 | sensory perception | 441 | -0.407502172 | -1.5517146 | 0.00023145 | 0.00023145 |
| GO:0034762 | regulation of transmembrane transport | 401 | -0.397706879 | -1.5037968 | 0.00023496 | 0.00023496 |
| GO:1904062 | regulation of cation transmembrane transport | 274 | -0.425667776 | -1.5640142 | 0.00024552 | 0.00024552 |
| GO:0007626 | locomotory behavior | 184 | -0.458616989 | -1.6245098 | 0.00025757 | 0.00025757 |
